# Extremely potent pan-sarbecovirus neutralizing antibodies generated by immunization of macaques with an AS03-adjuvanted monovalent subunit vaccine against SARS-CoV-2

**DOI:** 10.1101/2023.01.19.524784

**Authors:** Yupeng Feng, Meng Yuan, John M. Powers, Mengyun Hu, Jennifer E. Munt, Prabhu S. Arunachalam, Sarah R. Leist, Lorenza Bellusci, Lily E. Adams, Sumana Sundaramurthy, Lisa M. Shirreff, Michael L. Mallory, Trevor D. Scooby, Alberto Moreno, Derek T. O’Hagan, Harry Kleanthous, Francois J. Villinger, David Veesler, Neil P. King, Mehul S. Suthar, Surender Khurana, Ralph S. Baric, Ian A. Wilson, Bali Pulendran

## Abstract

The rapid emergence of SARS-CoV-2 variants that evade immunity to vaccination has placed a global health imperative on the development of therapeutic countermeasures that provide broad protection against SARS-CoV-2 and related sarbecoviruses. Here, we identified extremely potent pan-sarbecovirus antibodies from non-human primates vaccinated with an AS03 adjuvanted subunit vaccine against SARS-CoV-2 that recognize conserved epitopes in the receptor binding domain (RBD) with femtomolar affinities. Longitudinal analysis revealed progressive accumulation of somatic mutation in the immunoglobulin genes of antigen-specific memory B cells for at least one year following primary vaccination. 514 monoclonal antibodies (mAbs) were generated from antigen-specific memory B cells. Antibodies isolated at 5 to 12 months following vaccination displayed greater potency and breadth, relative to those identified at 1.4 months. Notably, 15 out of 338 (∼4.4%) antibodies isolated at 1.4∼6 months after the primary vaccination showed extraordinary neutralization potency against SARS-CoV-2 omicron BA.1, despite the absence of BA.1 neutralization in serum. Two of them, 25F9 and 20A7, neutralized authentic clade Ia sarbecoviruses (SARS-CoV, WIV-1, SHC014) and clade Ib sarbecoviruses (SARS-CoV-2 D614G, SARS-CoV-2 BA.1, Pangolin-GD) with half-maximal inhibition concentrations of (0.85 ng/ml, 3 ng/ml, 6 ng/ml, 6 ng/ml, 42 ng/ml, 6 ng/ml) and (13 ng/ml, 2 ng/ml, 18 ng/ml, 9 ng/ml, 6 ng/ml, 345 ng/ml), respectively. Furthermore, 20A7 and 27A12 showed potent neutralization against all SARS-CoV-2 variants of concern and multiple Omicron sublineages, including BA.1, BA.2, BA.3, BA.4/5, BQ.1, BQ.1.1 and XBB variants. X-ray crystallography studies revealed the molecular basis of broad and potent neutralization through targeting conserved RBD sites. In vivo prophylactic protection of 25F9, 20A7 and 27A12 was confirmed in aged Balb/c mice. Notably, administration of 25F9 provided complete protection against SARS-CoV-2, SARS-CoV-2 BA.1, SARS-CoV, and SHC014 challenge, underscoring that these mAbs are promising pan-sarbecovirus therapeutic antibodies.

**One Sentence Summary:** Extremely potent pan-sarbecovirus neutralizing antibodies

## INTRODUCTION

The zoonotic spillover of coronaviruses has caused three outbreaks of severe respiratory diseases within the last 20 years (*1-3*). SARS-CoV-2 has caused an enormous global health crisis, with over 650 million confirmed cases and over 6.6 million deaths globally as of December 22, 2022. Although COVID-19 vaccines and therapeutic antibodies have been developed at unprecedented speed, a series of variants of concern have emerged since late 2020. The antigenically distant B.1.1.529 (Omicron) variant first identified in Botswana in November 2021, which has spread worldwide with multiple sublineages that evade neutralization by antibodies has posed serious challenges to current vaccination strategies (*4-6*). Recent studies have revealed BNT162b2 vaccine (Pfizer/BioNTech) efficacy during the BA.4/5 waves was below 50% after two or three doses, with a fourth dose causing only a minimal, transient increase in the levels of Omicron-neutralizing antibodies (*4, 5*). Furthermore, currently available therapeutic antibodies suffer from a loss of efficacy against Omicron variants (*6, 7*), urging the need of broad protective vaccines (*8*) and monoclonal antibodies.

We recently performed a study in rhesus macaques to benchmark clinically relevant adjuvants, (including AS03, an α-tocopherol-containing oil-in-water emulsion; AS37, a Toll-like receptor 7 (TLR7) agonist adsorbed to alum; CpG1018-alum, a TLR9 agonist formulated in alum (CpG); Essai O/W 1849101, a squalene-in-water emulsion (OW); and Alum), for their capacity to enhance the protective immunity of SARS-CoV-2 vaccines, comprising either SARS-CoV-2 spike protein receptor-binding domain (RBD-NP) or a prefusion-stabilized spike (Hexapro-NP) on the surface of a self-assembling nanoparticle (*9*). We subsequently showed that a booster with RBD (beta)-NP, comprising the SARS-CoV-2 beta variant spike protein receptor-binding domain, at one year later elicited robust heterotypic protection against Omicron in macaques (*10*). Here, we analyzed banked samples from this study to investigate the evolution of the memory B cell response at the monoclonal level over a period of 1.5 years in rhesus macaques receiving the AS03 adjuvanted subunit vaccine. We observe significantly increased neutralization potency and breadth of mAbs isolated 6 to 12 months after the primary vaccination (the first two-dose vaccination) and 3 weeks to 6 months after the booster (the third dose vaccination at 12 months). Furthermore, we performed a serological assessment of the 15 most potent mAbs isolated 3 weeks to 6 months after the primary vaccination, performed a detailed structural analysis of a subset of these antibodies, and demonstrated their efficacy in protecting against Omicron variants of SARS-CoV-2 and other sarbecoviruses infections in mice.

## RESULTS

### Antibody and memory B cell responses

We analyzed the antigen-specific, memory B cell responses from banked PBMC samples from our previous study (*9*) in which rhesus macaques were immunized with RBD-NP adjuvanted with Alum, O/W, AS37, CpG, or AS03 (**fig. S1A**). We enumerated the circulating SARS-CoV-2 RBD specific IgG^+^ memory B cells (MBCs) by flow-cytometry using fluorescent-labeled probes (**fig. S1B**). The MBC frequency, variable (V) gene somatic hypermutation and CDR3 amino-acid length were comparable among differentially adjuvanted groups (**fig. S1, C to E**), suggesting that vaccination with these five different adjuvants exhibited a similar pattern of antigen-specific, memory B cell responses.

In addition, we analyzed memory B cell (MBC) responses using banked PBMC samples from our recent study (*10*) in which macaques were immunized with a two-dose primary vaccination of AS03-adjuvanted RBD-NP (N=5) or Hexapro-NP (N=6) at days 0 and 21, followed by a booster vaccination with AS03 adjuvanted RBD (beta)-NP one year later (**Fig. 1A**). We assessed the kinetics of the antibody response and MBCs in blood over 1.5 years in animals immunized with RBD-NP-AS03 or Hexapro-NP-AS03. Vaccination induced potent and broadly neutralizing antibodies against SARS-CoV-2 and Omicron variants after the booster (**Fig. 1b**). Interestingly, such Omicron neutralizing antibodies were not detectable at 5∼6 months following primary immunization, as described in our previous report (*10, 11*). We used flow cytometry to enumerate antigen-specific (RBD^+^ for RBD-NP group, Spike^+^ for Hexapro-NP group) MBCs in the blood (**Fig. 1C and fig. S1B**). The frequency of antigen-specific IgG^+^ MBCs peaked at 1.4 months and declined over the next 5∼6 months after vaccination (**Fig. 1D and fig. S2A**). Immunization with Hexapro-NP elicited a high proportion of IgG^+^ MBCs targeting the non-RBD region of spike protein, while there was no significant difference in RBD-specific IgG^+^ MBCs between RBD-NP and Hexapro-NP (**Fig. 1D**).

**Fig. 1.**
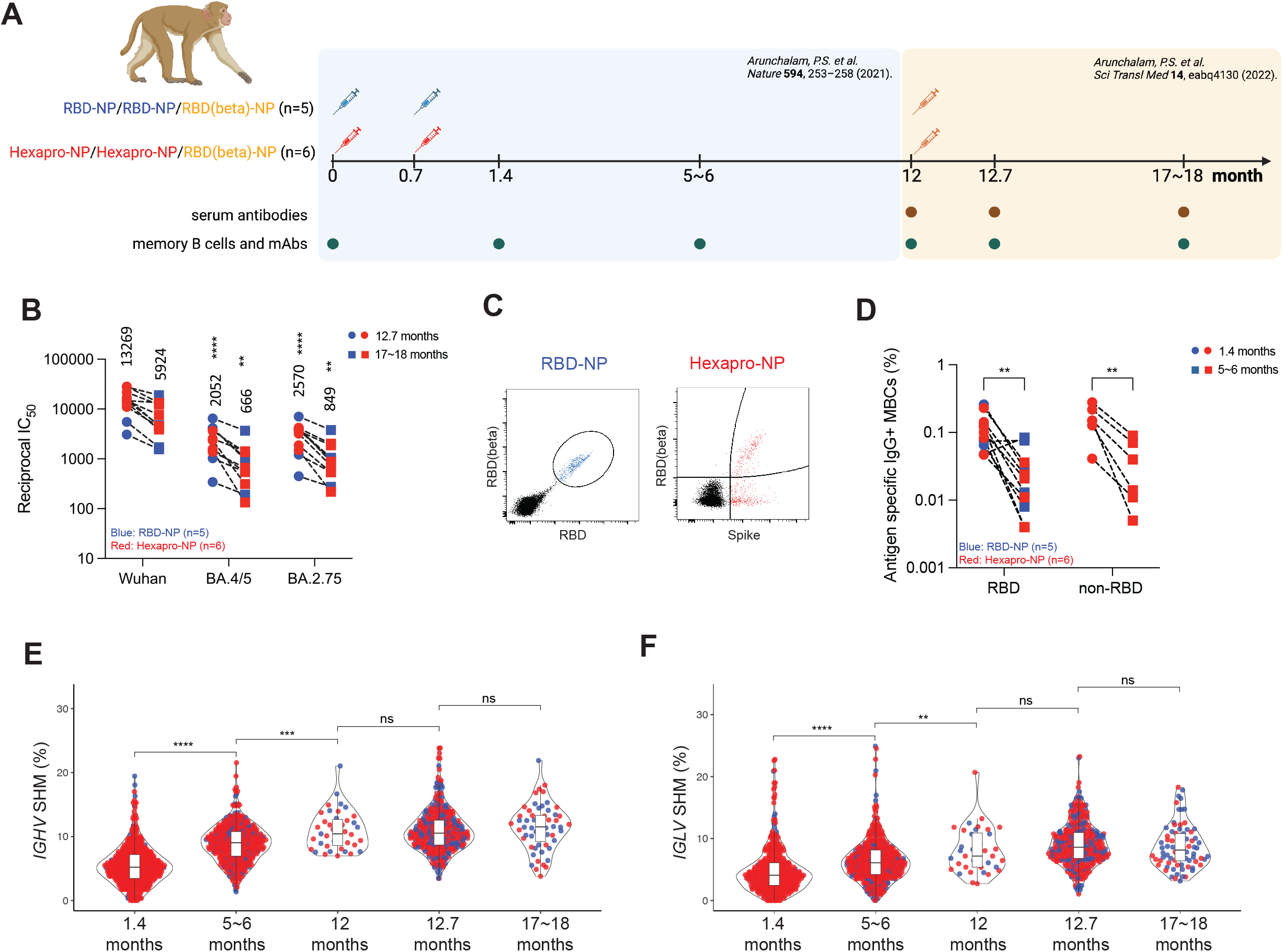
Antibody and memory B cell responses. (**A**) Study overview. Rhesus macaques received AS03 adjuvanted RBD-NP or Hexapro-NP on day 0 and day 21 and received a booster with RBD (beta)-NP with AS03 on 12 months. Blood samples were collected, and analysis was performed as illustrated in the diagram. (**B**) Pseudovirus neutralizing antibody responses against viruses indicated on X-axis are shown. Each symbol represents an animal [RBD-NP (blue; n = 5) and Hexapro-NP (red; n = 6)], and paired samples are connected with a dashed line. The numbers within the graphs show GMTs. The statistical differences were calculated using two-way ANOVA and the statistical differences between indicated viruses and SARS-CoV-2 Wuhan strain at the same time points were labeled as (**P < 0.01 and ****P < 0.0001). (**C**) Representative flow cytometry plots show dual RBD and RBD (beta) binding B cells for RBD-NP vaccinated animals (blue), and dual spike and RBD (beta) binding B cells for Hexapro-NP vaccinated animals (red). Cells were pre-gated on live, CD3^-^ CD14^-^ CD16^-^ CD20^+^ IgD^-^ IgM^-^ IgG^+^ B cells. (**D**) Frequency of antigen-specific IgG+ memory B cells relative to CD20^+^ B cells is shown for samples from the RBD-NP (blue) and Hexapro-NP (red) groups. The binding region is indicated on the X-axis. The statistical differences were calculated using two-way ANOVA (**P < 0.01). (**E**) Somatic hypermutation rates of the productive IGHV genes of B cells isolated from RBD-NP (blue) or Hexapro-NP (red) vaccinated animals at indicated time points. The boxes inside the violin plot show median, upper, and lower quartiles. Each dot represents an individual gene. The statistical differences between timepoints were calculated using one-way ANOVA (ns > 0.05, ***P < 0.001 and ****P < 0.0001). (**F**) as in (E), but for IGLV genes. The boxes inside the violin plot show median, upper, and lower quartiles. Each dot represents an individual gene. The statistical differences between timepoints were calculated using one-way ANOVA (ns > 0.05, **P < 0.01, and ****P < 0.0001).

Next, we analyzed somatic hypermutation (SHM) in the V genes of both heavy and light chains of antigen-specific, memory B cells, and observed a progressive increase in the frequency of SHM from 1.4 months to 5-6 months, plateauing at 12 months (**Fig. 1, E and F**). Interestingly, the booster immunization did not drive a further increase in somatic hypermutation (**Fig. 1, E and F**). To determine the germline gene usage of anti-SARS-CoV-2 antibodies in rhesus macaques, we compared V gene nucleotide sequences of SARS-CoV-2 spike-specific MBCs to the Macaca mulatta IG set from the IMGT database (*12*) and the Karolinska Macaque database (KIMDB) (*13*) using IgBLAST. Analyses showed that VH4-122 (IMGT) or VH4-93 (KIMDB) was most abundant (**fig. S2, B and C**). The closest germline gene of rhesus macaque VH4-122 in humans is VH4-59 (**fig. S2D**), a highly represented germline gene encoding anti-SARS-CoV-2 antibodies in humans (*13*). In humans, VH3-53 is one of the most frequently represented genes encoding antibodies generated in response to SARS-CoV-2 infection or mRNA vaccination. The structural basis of antibodies encoded by VH3-53 has been extensively studied and showed a highly convergent binding approach to the receptor binding site (*14-16*). The corresponding germline genes in rhesus macaque that are most similar to VH3-53 are VH3-103 (91.1%), VH3-100 (89.9%), and VH3S42 (89.9%), all of which have the SGGS motif in CDRH2, but no NY motif in CDRH1 (**fig. S2D**). Interestingly, we found those human VH3-53-like germline genes were also abundantly represented in rhesus macaques (**fig. S2B**).

### Isolation of monoclonal antibodies

To assess the evolution of the antibody repertoire encoded in antigen-specific, memory B cells, we sorted 3788 single SARS-CoV-2 Wuhan spike-specific IgG^+^ MBCs at indicated timepoints from macaques from all groups (RBD-NP plus all adjuvants and Hexapro-NP plus AS03) (**Fig. 1A and fig. S1A**), isolated 514 mAbs, and assessed their binding and neutralization potential (**Fig. 2A**). Consistent with the indistinguishable somatic hypermutation rates and CDR3 aa lengths (**fig. S1, D and E**), mAbs isolated from animals from the different adjuvanted groups displayed similar binding profiles (**fig. S3A**). Henceforth, all 514 mAbs were grouped per time for antibody evolution analysis.

**Fig. 2.**
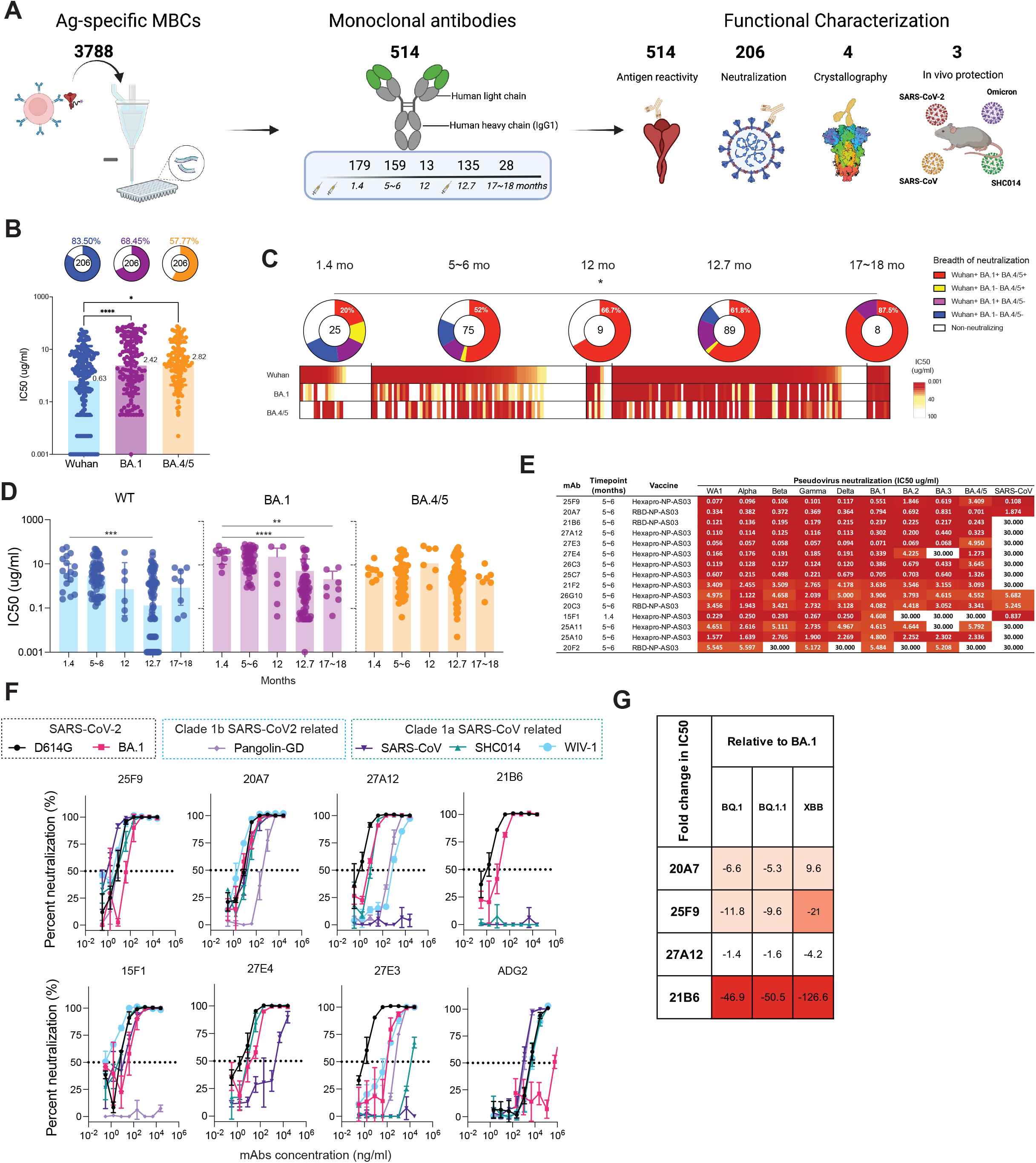
Characterization of monoclonal antibodies. (**A**) Diagram depicting the strategy for antigen-specific memory B cell sorting, monoclonal antibody isolation, and characterization. (**B**) Graphs showing the neutralizing activity of monoclonal antibodies measured by pseudovirion neutralization assay. Bar graphs show IC50 values for all neutralizing antibodies using SARS-CoV-2 Wuhan (blue), BA.1 (purple), and BA.4/5 (orange) pseudoviruses. Each dot represents one antibody. Pie charts illustrate the fraction of non-neutralizing (no valid IC50 or IC50 > 100 μg/ml) antibodies (white slices); the inner circles show the number of antibodies tested. The frequency of neutralizing antibodies against SARS-CoV-2 Wuhan (blue slices), BA.1 (purple slices), or BA.4/5 (orange slices) pseudoviruses was shown on top of each pie chart, respectively. Bars and whiskers indicate geometric mean and geometric SD. Statistical significance was determined by the two-tailed Kruskal–Wallis test with subsequent Dunn’s multiple-comparisons test (*P < 0.05 and ****P < 0.0001). (**C**) Heatmap shows the neutralization activity of mAbs isolated at indicated time points against pseudotyped SARS-CoV-2 Wuhan, BA.1, and BA.4/5 in rows respectively. Each unit within the heatmap represents one antibody. The color gradient indicates IC50 values ranging from 0 (red) to 100 (white). Pie charts illustrate the fraction of non-neutralizers (white slices), SARS-CoV-2 Wuhan only (blue slices), Wuhan and BA.1 double (purple slices), Wuhan and BA.4/5 double (orange slices), and Wuhan, BA.1, BA.4/5 triple (red slices) neutralizing antibodies; the inner circle shows the number of antibodies tested at indicated time points. Statistical significance between the frequencies of the five categories of antibodies isolated from five different time points was determined using a two-tailed chi-square test (*P < 0.05). (**D**) as in (B), graphs show kinetic change of the potency, reported as IC50 (μg/ml), of neutralizing antibodies against pseudotyped SARS-CoV-2 Wuhan (blue), BA.1 (orange) and BA.4/5 (purple), respectively. The statistical differences between timepoints were calculated using one-way ANOVA (**P < 0.01, ***P < 0.001 and ****P < 0.0001). (**E**) Heat maps showing the IC50 values of 15 selected monoclonal antibodies obtained from indicated vaccine group at indicated time points against the indicated pseudoviruses. The heat map range from 0.01 to 30 μg/ml is represented by white to dark red. f, Graphs show the neutralization of authentic SARS-CoV-2 D614, SARS-CoV-2 BA.1, Pangolin, SARS-CoV, SHC014, and WIV-1 by indicated antibodies. Symbols are means ± SD. Dashed lines indicate IC50 values. (**G**) The fold changes in neutralization IC50 values of BQ.1, BQ.1.1, XBB relative to BA.1, with resistance colored from white to dark red.

In total, 427 out of 514 (∼83.1%) mAbs bound to the SARS-CoV-2 Wuhan spike measured by enzyme-linked immunosorbent assay (ELISA), and the binding capacities (area under curve, AUC) strongly correlated with the V gene somatic hypermutation rates in the heavy chains (**fig. S3B**). This prompted us to examine whether B cell maturation over time drove affinity maturation. As expected, binding increased over time after the primary vaccination and plateaued at 12 months (**fig. S3C**). We further assayed the cross-reactivities of those mAbs against BA.1 and BA.4/5 by ELISA (**fig. S4A**). Interestingly, the correlation between Omicron-binding and somatic hypermutation was absent (**fig. S4, B to E**). However, the proportion of WT, BA.1, BA.4/5 triple-reactive mAbs increased over time and peaked at 12 months (**fig. S4F**).

Next, we selected the top 206 Omicron BA.1 binding mAbs for neutralization screening against pseudotyped SARS-CoV-2 Wuhan, BA.1, and BA.4/5 strains. 83.5% of the 206 mAbs neutralized SARS-CoV-2 Wuhan, while 68.5% and 57.8% neutralized Omicron BA.1 and BA.4/5, respectively [**Fig. 2B (top)**]. The average potency of neutralizing antibodies against BA.1 or BA.4/5 was also significantly lower than against Wuhan strain, reflecting the fact that the vaccine contained the Wuhan strain [**Fig. 2B (bottom)**]. To determine whether antibody maturation in binding and cross-reactivity translated into enhanced neutralizing potency and breadth, we analyzed the neutralization profiles as a function of time post-immunization.

Remarkably, there was a significant increase in the frequency of SARS-CoV-2 Wuhan, BA.1, and BA.4/5 triple-neutralizing antibodies, and a decrease in the percentage of non-neutralizing antibodies over 1.5 years, an indication of the evolution in the neutralization potency and breadth (**Fig. 2C**). Further examining the consequence of this evolution, we found that the average potency of neutralization against SARS-CoV-2 Wuhan and BA.1 was significantly improved by the booster immunization (**Fig. 2D**). Collectively, our data demonstrate an evolution of monoclonal antibodies in the memory B cell compartment towards higher neutralization potency and breadth.

### Potent broadly neutralizing antibodies

To identify SARS-CoV-2 neutralizing antibodies with both high potency and pan-sarbecovirus breadth, we further characterized the neutralizing antibodies isolated at 1.4 months and 5∼6 months and identified 15 mAbs showing better or comparable neutralizing activity against the SARS-CoV-2 BA.1 variant in a side-by-side comparison assay with a recently described Omicron BA.1 neutralizing antibody S2H97 (*17*) (**fig. S5A**). All 15 mAbs displayed high avidities [apparent K_D_ < 0.1 ng/ml (∼666 femtomolar)] against RBDs of different SARS-CoV-2 variants of concern measured by biolayer interferometry binding assay (**Table 1**), and 8 mAbs showed strong avidity (K_D_: 0.015∼12.4 nM) to the spike of SARS-CoV (**Table 2**). The potency and breadth of these 15 mAbs were further confirmed by neutralization of a panel of 10 pseudoviruses carrying spikes of SARS-CoV-2 WA1, 8 SARS-CoV-2 variants of concern (alpha, beta, gamma, delta, BA.1, BA.2, BA.3, BA.4/5), and SARS-CoV (fig. S5B). Five bnAbs (25F9, 20A7, 21B6, 27A12, 27E3) stood out for their high potency and breadth across the SARS-CoV-2 variants (**Fig. 2E**). Remarkably, 20A7 can neutralize all SARS-CoV-2 Omicron variants and SARS-CoV with little to no reduced potency as compared to SARS-CoV-2 WA1 strain (**Fig. 2E**).

**Table 1.** Binding affinities of indicated monoclonal antibodies against RBDs of SARS-CoV-2 WT, Omicron BA.1, Alpha, Beta and Delta variants.

**RBD Binding** (Apparent KD, ng/ml)
|  | WT | Omicron | Alpha | Beta | Delta |
| --- | --- | --- | --- | --- | --- |
| 27A12 | <0.1 | 0.3 | <0.1 | <0.1 | <0.1 |
| 27E3 | <0.1 | 14.4 | <0.1 | 18.9 | <0.1 |
| 27E4 | <0.1 | <0.1 | <0.1 | <0.1 | <0.1 |
| 21B6 | <0.1 | <0.1 | <0.1 | <0.1 | 4.1 |
| 26G10 | <0.1 | <0.1 | <0.1 | <0.1 | <0.1 |
| 20A7 | <0.1 | <0.1 | <0.1 | <0.1 | <0.1 |
| 25F9 | <0.1 | <0.1 | <0.1 | <0.1 | <0.1 |
| 15F1 | 1.0 | <0.1 | <0.1 | <0.1 | <0.1 |
| 26C3 | <0.1 | 11.4 | <0.1 | 2.0 | <0.1 |
| 25C7 | <0.1 | <0.1 | <0.1 | <0.1 | <0.1 |
| 21F2 | <0.1 | <0.1 | <0.1 | <0.1 | <0.1 |
| 25A11 | <0.1 | <0.1 | <0.1 | <0.1 | <0.1 |
| 20F2 | 0.1 | <0.1 | <0.1 | <0.1 | <0.1 |
| 25A10 | <0.1 | <0.1 | <0.1 | <0.1 | <0.1 |
| 20C3 | <0.1 | <0.1 | <0.1 | <0.1 | <0.1 |
| CR3022 | 14.1 | 17.4 | 12.0 | 12.7 | 8.3 |
| CC12.3 | 5.8 | n.b. | n.a. | n.a. | n.a. |

n.b. = no binding
n.a. = data not available

**Table 2.**
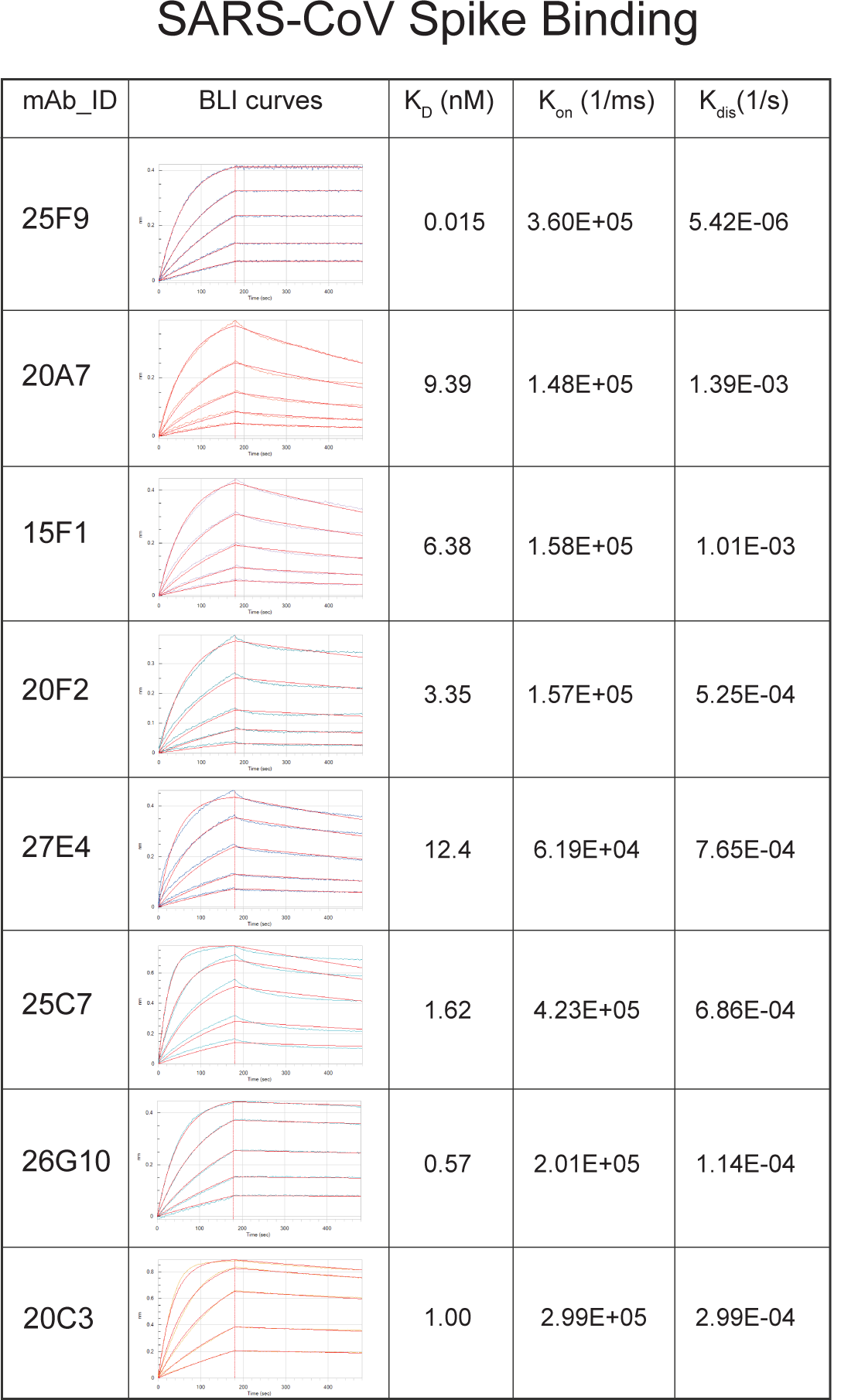
Binding affinities of indicated monoclonal antibodies against Spikes of SARS-CoV.

SARS-CoV Spike Binding
| mAb_ID | BLI curves | $K_D$ (nM) | $K_{on}$ (1/ms) | $K_{dis}$ (1/s) |
| --- | --- | --- | --- | --- |
| 25F9 | | 0.015 | $3.60E+05$ | $5.42E-06$ |
| 20A7 | | 9.39 | $1.48E+05$ | $1.39E-03$ |
| 15F1 | | 6.38 | $1.58E+05$ | $1.01E-03$ |
| 20F2 | | 3.35 | $1.57E+05$ | $5.25E-04$ |
| 27E4 | | 12.4 | $6.19E+04$ | $7.65E-04$ |
| 25C7 | | 1.62 | $4.23E+05$ | $6.86E-04$ |
| 26G10 | | 0.57 | $2.01E+05$ | $1.14E-04$ |
| 20C3 | | 1.00 | $2.99E+05$ | $2.99E-04$ |

We next evaluated the neutralization of authentic SARS-CoV-2 D614G, SARS-CoV-2 BA.1, GD Pangolin-CoV, SARS-CoV, SHC014, WIV-1, and MERS-CoV by 7 bnAbs (25F9, 20A7, 21B6, 27A12, 27E3, 27E4, 15F1) and a previously described ultrapotent neutralizing antibody ADG2 (*18*). Strikingly, 25F9 neutralized all the above SARS-related viruses with IC50 values of 6 ng/ml, 42 ng/ml, 6 ng/ml, 0.85 ng/ml, 3 ng/ml, 6 ng/ml, respectively, with potencies surpassing that observed with ADG2 in a head-to-head comparison in our assay (**Fig. 2F, and fig. S5C**).

20A7 displayed similar neutralizing breadth as compared to 25F9, albeit there was a reduction of neutralization against Pangolin, whereas none neutralized MERS-CoV due to its distinct RBD (**fig. S5C**). Of note, two SARS-CoV-2 strain-specific bnAbs, 21B6 and 27A12, neutralized SARS-CoV-2 BA.1 with IC50 values of 11 ng/ml and 5 ng/ml, respectively, consistent with the above pseudovirus neutralization data (**Fig. 2D and fig. S5C**). During the conduction of the study, new variants of concern, such as BQ.1.1 and XBB, become the dominating circulating viruses. We assayed 25F9, 20A7, 27A12, and 21B6 for their neutralization against BQ.1, BQ.1.1 and XBB using the same pseudovirus neutralization approach (**Fig. 2G, and fig. S5D**).

Interestingly, 27A12 exhibited comparable neutralization potencies against BQ.1, BQ.1.1 and XBB compared to that against BA.1 (**Fig. 2G**). 20A7 and 25F9 showed some reduction (**Fig. 2G**). However, their extremely high authentic BA.1 neutralization potencies (6 ng/ml and 42 ng/ml, respectively) (**Fig. 2F**) make 20A7 and 25F9 still quite competitive to recently described antibodies (*7*). Altogether our data revealed the cellular and molecular basis of vaccine-induced antibody evolution. More importantly, we isolated some extremely potent SARS-CoV-2 strain-specific bnAbs (21B6 and 27A12) and potent pan-sarbecovirus bnAbs (25F9 and 20A7) after the primary vaccination with a nanoparticle immunogen adjuvanted with AS03.

### Structural basis of neutralization breadth

To define the binding epitopes of the bnAbs and the structural basis of their neutralization breadth, we performed competitive binding experiments using biotinylated antibodies (CR3022, CC12.3, CV07-270, S2X259, S2M11), whose epitopes are well characterized (*14, 19-22*). Interestingly, the top three potent Omicron neutralizers (27A12, 27E3, 21B6) competed with the ultrapotent monoclonal antibody S2M11 (*22*), which recognizes epitopes overlapping with the RBM (**fig. S6, A and B**). Co-incubation of the top four antibodies with the greatest neutralization breadth (25F9, 20A7, 15F1, 27E4) with CR3022 (*14*), a SARS-CoV neutralizing antibody, or with S2X259 (*21*), a pan-sarbecovirus neutralizing antibody, showed very strong competition (83-96%), suggesting some similarity in their binding epitopes (**fig. S6, A and B**).

Next, we applied X-ray crystallography to determine the crystal structures of SARS-CoV-2 RBD in complex with three antibodies isolated in this study (**Table S3**), 25F9 (**Fig. 3**), 20A7 (**Fig. 4**), and 21B6 (**fig. S7**). Relative positions of epitopes of the three antibodies as well as the receptor binding site (RBS), the CR3022 site, and the S309 (23) site are shown in **Fig. 3A**. 25F9 targets one side of the RBD with some overlap with the conserved CR3022 site (**Fig. 3B**), where approximately 80% of the 25F9 epitope is buried by the heavy chain (**Fig. 3C**). CDRs H2, H3, L1, L3, and LFR3 interact with RBD (**Fig. 3D**). 25F9 binding would clash with the human receptor ACE2 (**Fig. 3E**), which explains its high neutralization potency. 25F9 targets a conserved region of the RBD, where 23-27 out of 28 epitope residues are conserved among SARS-CoV-2 variants, including BQ.1.1 and XBB.1.5, and other SARS-like viruses, e.g. SARS-CoV (SARS1), pang17, and RaTG13 (**Fig. 3F**), which further explains the high neutralization potency of 25F9 against a broad range of SARS-like viruses (**Fig. 2, E and F**). 25F9 V_H_ F54 inserts into a hydrophobic pocket in the RBD and stacks with aromatic residues RBD-Y365, F377, Y369, P384, and aliphatic residue L387 (**Fig. 3G**). 25F9-V_H_ K52 side chain and V_H_ A52c backbone carbonyl hydrogen bond (HB) with RBD-Y369 side-chain hydroxyl and C379 backbone amide, respectively (**Fig. 3G**). The CDR H3 and light chain of 25F9 also form polar and hydrophobic interactions with the RBD (**Fig. 3, H and I**).

**Fig. 3.**
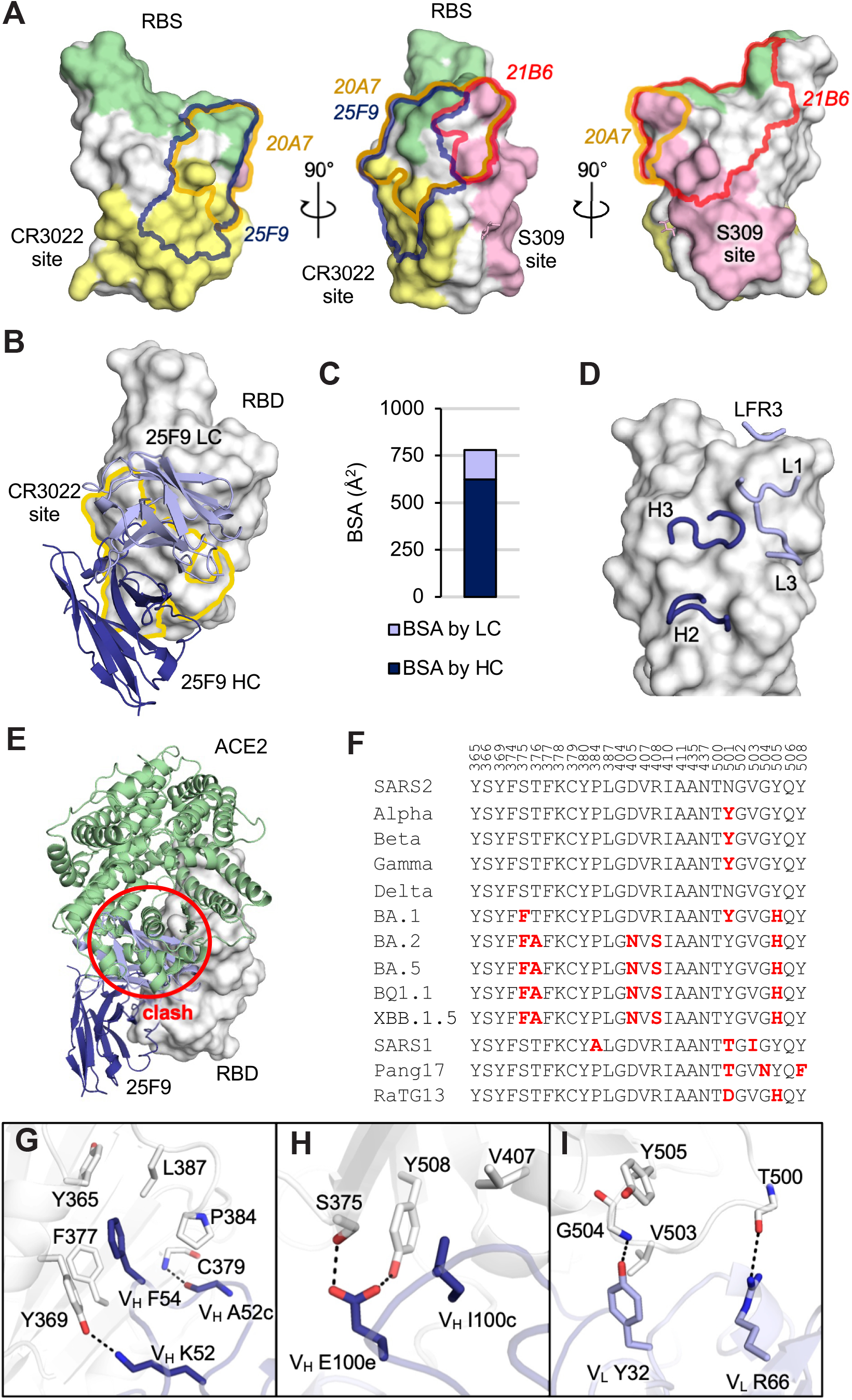
Crystal structure of 25F9 in complex with SARS-CoV-2 RBD. The SARS-CoV-2 RBD is shown in white and human ACE2 (hACE2) is in pale green throughout all the figures, while heavy and light chains of 25F9 are in blue and lavender. For clarity, only variable domains of the antibodies are shown in all figures. (**A**) Relative positions of epitopes on SARS-CoV-2 RBD (white). The receptor binding site (RBS) is shown in pale green, CR3022 site in yellow, S309 site in pink, with epitopes of 25F9, 20A7, and 21B6 highlighted in blue, orange, and red outlines, respectively. RBS and epitope residues are defined as buried surface area (BSA > 0 Å^2^) as calculated by Proteins, Interfaces, Structures and Assemblies (PISA, http://www.ebi.ac.uk/pdbe/prot_int/pistart.html). (**B**) Crystal structure of 25F9 in complex with SARS-CoV-2 RBD. (**C**) Surface area of SARS-CoV-2 buried by heavy and light chains of 25F9. (**D**) 25F9 interacts with RBD using CDRs H2, H3, L1, L3, and LFR3. (**E**) SARS-CoV-2 RBD with 25F9 superimposed onto an RBD/hACE2 complex structure (PDB 6M0J) shows that 25F9 would clash (indicated with a red circle) with hACE2 receptor (**F**) Sequence alignment of epitope residues in a subset of SARS-like viruses. Residues that differ from wild-type SARS-CoV-2 are indicated in red. (**G-I**) Molecular interactions between RBD and (G) CDR H2, (H) CDR H3, and (I) light chain. Hydrogen bonds and salt bridges are indicated by dashed lines.

**Fig. 4.**
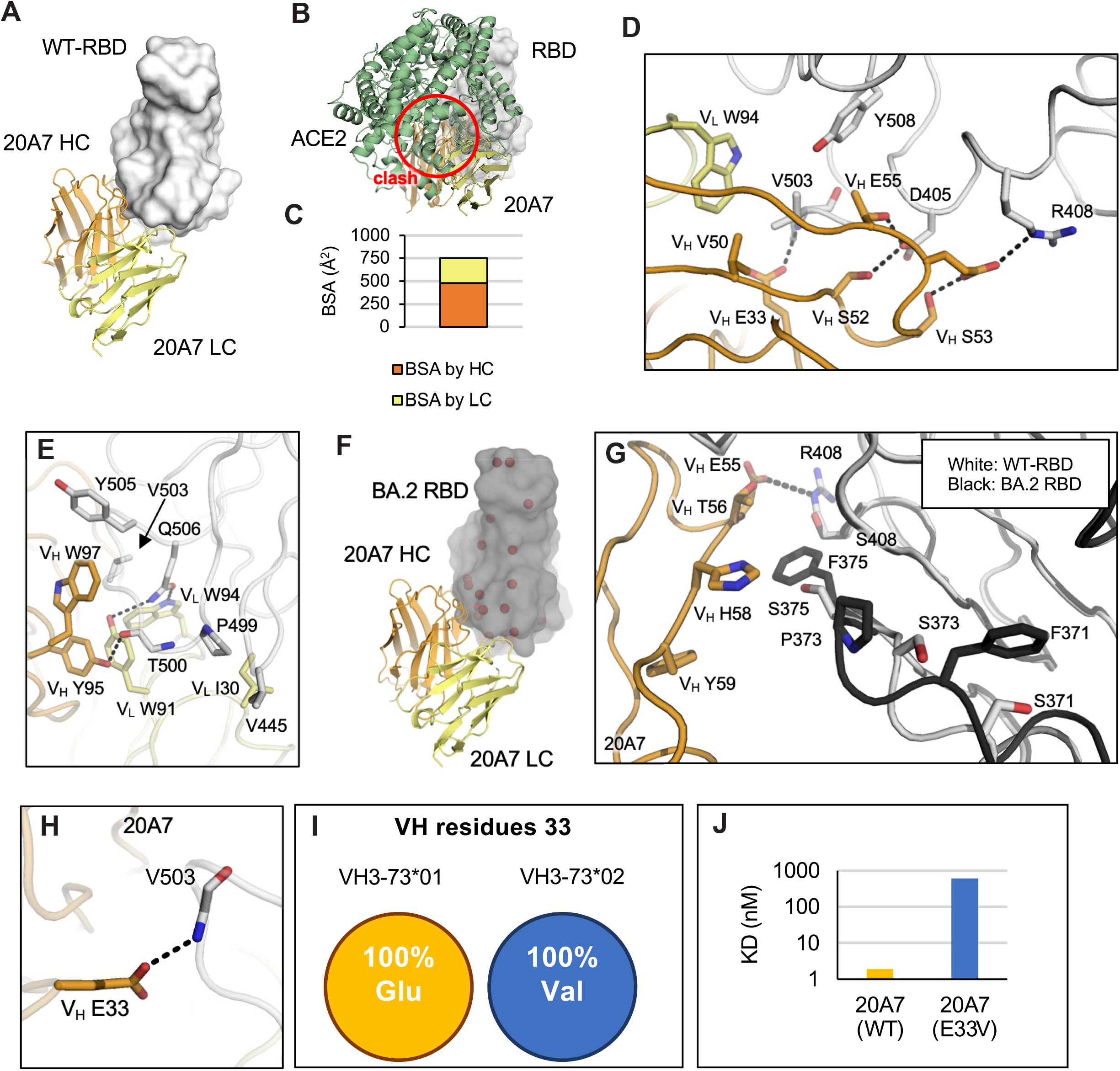
Crystal structures of 20A7 in complex with SARS-CoV-2 wild-type and BA.2 RBDs. The SARS-CoV-2 wild-type and BA.2 RBDs are shown in white and black, while the heavy and light chains of 20A7 are in orange and yellow. Hydrogen bonds and salt bridges are indicated by dashed lines. (**A**) Crystal structure of 20A7 in complex with SARS-CoV-2 wild-type RBD. (**B**) SARS-CoV-2 RBD in complex with 20A7 superimposed onto an RBD/hACE2 complex structure (PDB 6M0J) shows that 20A7 would clash (red circle) with the hACE2 receptor (**C**) Surface area of SARS-CoV-2 wild-type RBD buried by heavy and light chains of 20A7. (**D-E**) Molecular interactions between wild-type SARS-CoV-2 RBD and 21B6. (**F**) The crystal structure of 20A7 with RBD (BA.2) shows that 20A7 targets BA.2 in the same binding mode as wild-type SARS-CoV-2. Mutated residues in the Omicron BA.2 subvariant are indicated by red spheres. (**G**) Structural comparison of the interaction of 20A7 with wild-type (white) and BA.2 (black) RBDs. (**H**) 20A7 V_H_ E33 forms a hydrogen bond with RBD. (**I**) All alleles of VH3-73*01 encode Glu at position 33 whereas those of VH3-73*02 encode Val. (**J**) BLI binding assay showed that E33V reduced the binding of 20A7 to SARS-CoV-2 RBD by ∼300 fold.

21B6 neutralizes a broad range of SARS-CoV-2 variants, including BA.2 and BA.5, with high potency but does not neutralize other SARS-like viruses SARS-CoV and SHC014 (**Fig. 2, E and F**). In contrast, 21B6 binds the opposite side (above the S309 epitope) of SARS-CoV-2 RBD (**fig. S7A**). Like 25F9, 21B6 targets RBD with approximately 80% of its epitope area buried by the heavy chain (**fig. S7B**), and CDRs H1, H2, H3, L2, and HFR1 interact with the RBD (**fig. S7C**). 21B6 would also clash with hACE2 (**fig. S7D**). The 21B6 epitope residues are conserved from wild-type SARS-CoV-2 through Omicron subvariants BA.2 and BA.5, but vary in other SARS-like viruses (**fig. S7E**). RBD-Y351 hydrogen bonds with 21B6-V_H_ S30 and D31 and makes hydrophobic interactions with V_H_ T28 and Y32 (**fig. S7F**). CDR H2, especially V_H_ V52b and L52c, extensively interact with a hydrophobic patch on RBD formed by L452, Y351, L492, F490, and T470 (**fig. S7G**). CDR H3 and the light chain of 21B6 also form extensive interactions with the RBD (**fig. S7, H and I**). Notably, the R346T mutation in the most recent circulating Omicron subvariants BQ.1.1 and XBB.1.5 would cause a loss of salt bridge with 21B6-V_H_ D101, which may contribute to some loss of neutralization to the new subvariants (**Fig. 2G**).

20A7 neutralizes all SARS-CoV-2 variants as well as SARS-CoV (**Fig. 2, E and F**). The crystal structure of 20A7 with wild-type SARS-CoV-2 RBD (**Fig. 4A**) shows its binding to RBD would clash with ACE2 (**Fig. 4B**). Both heavy and light chains interact with the RBD, where the relative RBD surface area buried by heavy and light chains is approximately 2/3 and 1/3, respectively (**Fig. 4C**). CDRs H2 (**Fig. 4D**) and H3 (**Fig. 4E**) form extensive interactions with the RBD. 20A7 neutralizes Omicron subvariants with little reduction in activity (**Fig. 2, E and F**). We also determined a crystal structure of 20A7 in complex with Omicron BA.2 RBD (**Fig. 4F**) that exhibited the same binding mode as with wild-type SARS-CoV-2 RBD (**Fig. 4A**).

Comparison of the interactions of 20A7 with wild-type and BA.2 variant (**Fig. 4G**) showed that the salt bridge formed by 20A7 V_H_ E55 and RBD-R408 was eliminated by the R408S mutation in BA.2, which may contribute to some loss of binding affinity. In addition, mutations at S371, S373, and S375 in the Omicron subvariants induce a localized conformational change in the main chain of a loop containing these residues in the RBD (24).

Some of the authors of this study previously determined two antibody structures encoded by a highly enriched germline gene VH3-73 in SARS-CoV-2-immunized macaques (*25*). In the present study, we show that another VH3-73-encoded antibody, 20A7, adopts the same binding mode (**fig. S8, A to C**). Importantly, V_H_ E33 in all three antibodies hydrogen bonds with the backbone amide of RBD-V503. V_H_ E33 is only encoded by the alleles clustered as VH3-73*01, whereas VH3-73*02 alleles encode a valine instead of glutamic acid, where a single mutation 20A7-V_H_ E33V markedly reduced the binding affinity by ∼300 fold, which was presumably due to the loss of this HB (**Fig. 4, H to J, and fig. S8**), suggesting that it may be difficult to generate such broadly neutralizing antibodies in macaques that do not have an VH3-73*01 allele. We previously made a similar observation that human anti-SARS-CoV-2 VH2-5/VL2-14 antibodies are also allele-specific (*26*). These observations highlight the importance of considering allele polymorphism in vaccine design.

We previously discovered a broad-and-potent neutralization epitope on the RBDs of SARS-related coronaviruses (*27*). The epitope spans from a corner of RBS-D to the CR3022 site (**fig. S9**). Here we show that the broad-and-potent neutralizing antibodies 25F9 and 20A7 also target this RBS-D/CR3022 site and neutralize all tested SARS-CoV-2 variants as well as other SARS-like viruses. Furthermore, a potent neutralizing antibody SA55 in a clinical trial potently neutralizes all known SARS-CoV-2 variants including BQ.1.1 and XBB and also targets the RBS-D/CR3022 site (*28*) (**fig. S9**). The features of this site that elicit recurring broad-and-potent neutralizing antibodies provide a promising target for universal COVID-19 vaccines.

### Broad sarbecovirus protection in mice

To evaluate the broad protection of the best two pan-sarbecovirus bnAbs (25F9 and 20A7) and the best Omicron-neutralizing bnAb (27A12), we conducted prophylactic challenge studies in 12 months old female BALB/c mice with mouse-adapted (MA) sarbecoviruses, including SARS-CoV-2 (MA10), SARS-CoV-2 BA.1, and SARS-CoV (MA15) (*29-31*). We conducted an additional SHC014 challenge study (*32*) for 25F9 because of its extensive neutralization breadth and potency (**Fig. 5A**). The bnAbs were administered by intraperitoneal injection at 200 μg per mouse 12 hours before intranasal administration of viruses with 10^3^ plaque-forming units (PFU) for SARS-CoV-2 (MA10), 10^4^ PFU for SARS-CoV (MA15), and 10^5^ PFU for SARS-CoV-2 BA.1 and SHC014 (MA15). Mice were monitored for daily weight changes and lung tissues were collected two- or four-days post-infection (dpi) for gross pathology assessment, virus quantification, histology, and immunohistochemistry analysis. Mice treated with isotype control antibody exhibited substantial and progressive weight loss due to the infection of all viruses.

**Fig. 5.**
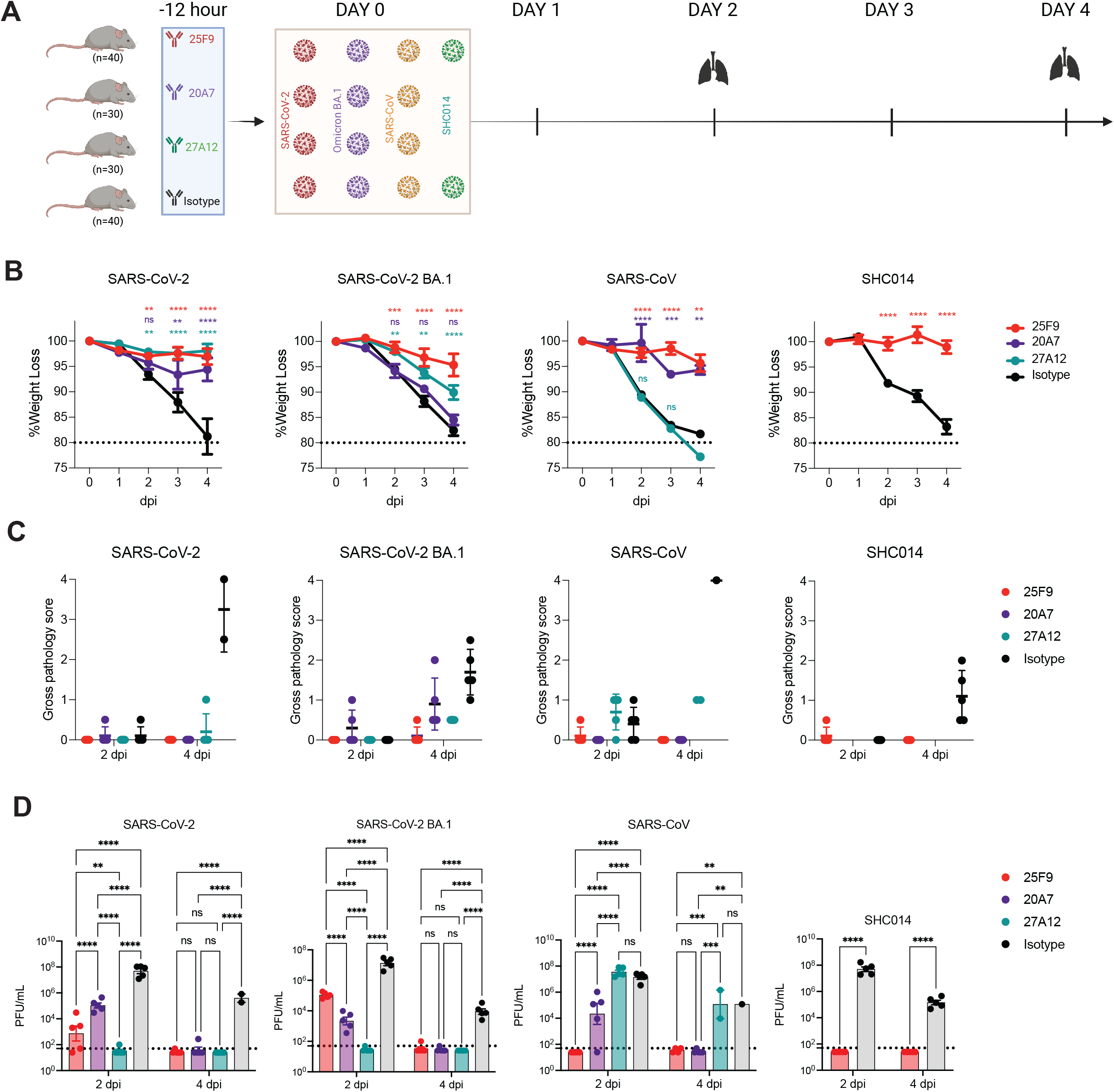
25F9, 20A7, and 27A12 protect aged mice from sarbecovirus related diseases. (**A**) Diagram depicting the challenge study in mice. 25F9, 20A7, 27A12, or a DENV2(2D22) control antibody were administered intraperitoneally at 200 μg per animal into 14 groups of aged mice (10 animals per group). Each group of animals was challenged intranasally 12 h after antibody infusion with one of the indicated sarbecoviruses (mouse-adapted SARS-CoV-2, 1 × 10^3^ PFU; mouse-adapted SARS-CoV-2-BA.1 chimera, 1 × 10^5^ PFU; mouse-adapted SARS-CoV, 1 × 10^4^ PFU; or SHC014 MA15 chimera, 1 × 10^5^ PFU). As a control, groups of mice were exposed only to PBS in the absence of virus. (**B**) The body weight change of mice after challenge with moused-adapted SARS-CoV2, SARS-CoV-2 BA.1, SARS-CoV and SHC014, respectively (mean ± SEM from 10 animals per group from days 0 to 2, n = 5 from days 3 to 4; mixed-effects model with post hoc Dunnett’s multiple tests in comparison with the control group; significance levels are shown as **p < 0.01, ***p < 0.001, and ****p < 0.0001 or ns when not significant. (**C**) Lung gross pathology was scored at the collection on day 2 and 4 post-infection in mice prophylactically treated with indicated bnAbs or the isotype control mAb (n = 5 per group). Data are presented as mean values ± SEM. (**D**) Lung virus titers (PFU per lung) were determined by plaque assay of lung tissues collected at days 2 or 4 after infection (n = 5 individuals per time point for each group). Data are shown as scatter dot plots with bar heights representing the mean and whiskers representing SEM. mixed-effects model with post hoc Dunnett’s multiple tests in comparison with the control group; significance levels are shown as **p < 0.01, ***p < 0.001, and ****p < 0.0001 or ns when not significant.

25F9 completely prevented weight loss from SARS-CoV-2 (MA10), SARS-CoV (MA15), and SHC014 (MA15) infection (**Fig. 5B**). Interestingly, although 20A7 and 27A12 showed much higher neutralization of SARS-CoV-2 BA.1 than that of 25F9 (IC50: 6 ng/ml, 5 ng/ml, and 42 ng/ml, respectively) in vitro (**fig. S5C**), 25F9 showed the best protection from weight loss after SARS-CoV-2 BA.1 infection (**Fig. 5B**). No signs of lung discoloration (gross pathology) were observed at either 2 or 4 dpi in mice treated with 25F9 (Fig. 5c). Furthermore, we assessed the viral load in the lungs. 25F9 completely abrogated viral replication in all mice at 4 dpi (**Fig. 5D**). Consistent with *in vitro* neutralization, 20A7 and 27A12 prophylactic treatment leads to better BA.1 clearance in vivo than 25F9 at 2 dpi (**Fig. 5D**), suggesting 25F9 potentially utilized multi-function for disease protection. We conclude that all three mAbs effectively protected against SARS-CoV-2 (MA10) infection in mice. 25F9 protected against the SARS-CoV-2 Omicron variant and other sarbecoviruses equally effectively as it protected against the SARS-CoV-2 ancestral strain, highlighting its potential as a pan-sarbecovirus therapeutic antibody.

## DISCUSSION

Memory B cells, a fundamental component of human immunity to SARS-CoV-2, mature over time after natural infection or mRNA vaccination (*33-37*). We recently showed that an AS03, a squalene oil-in-water emulsion adjuvant developed by GlaxoSmithKline, adjuvanted nanoparticle vaccine conferred durable and heterotypic protection against Omicron challenge with 100% and ∼65% protection at 6 weeks and 6 months post the booster, respectively (*10*). The rapid elicitation of broad and potent serum neutralizing antibodies following the booster immunization, suggested the evolution of a broad and potent antibody repertoire encoded in the memory B cell compartment. Consistent with this notion, we found in this study that somatic mutations and the potency and breadth of antibodies encoded by B cell receptors in memory B cells evolved after the primary vaccination. Those matured memory B cells with greater potency and breadth can rapidly differentiate into antibody-secreting cells in response to a booster immunization or infection. Although it is well-known that adjuvants can modulate and enhance the magnitude, breadth, and durability of the vaccine-induced serum antibody response, few studies have investigated their effects on the monoclonal level (*26, 38-40*). In this study, we found the primary vaccination of the AS03-adjuvanted nanoparticle-based subunit vaccine elicited a progressive antibody evolution towards greater potency and breadth over a period of one year, presumably driven by antigen-antibody complexes on follicular dendritic cells. Whether the same level of antibody maturation and bnAbs will be observed without adjuvant or in humans needs further investigation.

Due to the scarcity of effective mAbs against the Omicron variants (*6, 7, 41-43*) and the potential for zoonotic coronaviruses such as SARS-CoV, MERS-CoV as well as bat coronaviruses WIV-1, RaTG13, SHC014, etc. (*2, 3, 31, 44-48*) to spill-over into humans, considerable effort is focused on identifying broad neutralizing antibodies able to cross-neutralize various SARS-CoV-2 variants and other SARS-related viruses. However, there is always a compromise between potency and breadth (*16, 18, 41, 49-51*). Here we identified 7 bnAbs showing potent neutralization against authentic SARS-CoV-2 WA1 strain with IC50s below 10 ng/ml. All these 7 mAbs neutralized previous SARS-CoV-2 variants of concern without any reduction in potency. 25F9, 20A7, 21B6, and 27A12 neutralized authentic SARS-CoV-2 BA.1 with IC50s of 42 ng/ml, 6 ng/ml, 11 ng/ml, 5 ng/ml, respectively. Notably, 27A12 showed little if any reduction in their neutralization against SARS-CoV-2 BA.2, BA.3, BA.4/5, BQ.1, BQ.1.1 and XBB relative to SARS-CoV-2 BA.1. 25F9 and 20A7 neutralized authentic SARS-CoV and several other bat coronaviruses with comparable potency as compared to that against SARS-CoV-2, albeit 20A7 showed some reduction of neutralization against a Pangolin strain. Furthermore, we determined crystal structures of mAbs (25F9, 20A7, and 21B6) and their mode of binding to the RBD of SARS-CoV-2 and one structure of 21B6 complexed with the RBD of SARS-CoV-2 BA.2 at resolutions of 3.05 Å, 2.58 Å, 1.75 Å and 2.30 Å, respectively.

Finally, the most potent and broadly neutralizing antibodies, 25F9, 20A7, and 27A12, were evaluated for their prophylactic protection efficacy against four different SARS-related viruses, including mouse-adapted SARS-CoV-2 MA10, SARS-CoV-2 BA.1, SARS-CoV MA15 and SHC014 MA15 in aged mice. This mouse model is well recognized for its efficient recapitulation of disease pathogenesis (*52*). Our results demonstrated that prophylactic treatment with a single dose of bnAbs not only led to clinical improvement, as shown by the absence of weight loss, but also to markedly reduced lung pathology, virus load and inflammatory infiltration. In conclusion, the identification of two ultrapotent broadly neutralizing antibodies, 25F9 and 20A7, provides, to our knowledge, two of the most potent and broadest neutralizing antibodies described to date, making them promising candidates for treating sarbecovirus infection.

## List of Supplementary Materials

Materials and Methods

Fig. S1 to S9

Table S1 to S3

Data file S1

References (*53–65*)

## Acknowledgments

We thank the staff at the NIRC, UL Lafayette for conducting the animal experiment and sample collection; the Stanford FACS facility for maintenance and access to flow cytometers and FACS sorting; and all the members of GlaxoSmithKline (GSK) for critical reading of the manuscript. We are grateful to the staff of Advanced Photon Source and Stanford Synchrotron Radiation Lightsource (SSRL) Beamline 12-1 for assistance. GM/CA@APS has been funded by the National Cancer Institute (ACB-12002) and the National Institute of General Medical Sciences (AGM-12006, P30GM138396). This research used resources of the Advanced Photon Source; a U.S. Department of Energy (DOE) Office of Science User Facility operated for the DOE Office of Science by Argonne National Laboratory under Contract No. DE-AC02-06CH11357. Extraordinary facility operations were supported in part by the DOE Office of Science through the National Virtual Biotechnology Laboratory, a consortium of DOE national laboratories focused on the response to COVID-19, with funding provided by the Coronavirus CARES Act. Use of the Stanford Synchrotron Radiation Lightsource, SLAC National Accelerator Laboratory, is supported by the U.S. Department of Energy, Office of Science, Office of Basic Energy Sciences under Contract No. DE-AC02-76SF00515. The SSRL Structural Molecular Biology Program is supported by the DOE Office of Biological and Environmental Research, and by the National Institutes of Health, National Institute of General Medical Sciences (P30GM133894). The UNC Animal Histopathology & Laboratory Medicine Core is supported in part by an NCI Center Core Support Grant (5P30CA016086-41) to the UNC Lineberger Comprehensive Cancer Center. Cartoons were created with BioRender.com.

## Funding

This study was supported by the Bill and Melinda Gates Foundation INV-018675 (to BP) and INV-004923 (to IAW) and NIH PO1 AI167966 (RSB) and INV-010680 (to D.V. and N.P.K.); the National Institutes of Health (1P01AI167966 to F.J.V., D.V., N.P.K., R.S.B., and B.P.)

## Author contributions

Conceptualization: BP and YF

Investigation: YF, MY, JMP, MH, JEM, PSA, SRL, LB, LEA, SS, LMS, MLM, TDS, and AM

Data curation and analysis: YF, MY, JMP, and JEM

Visualization: YF and MY

Funding acquisition: BP, IAW, and RSB

Project administration: BP and YF

Supervision: BP, IAW, RSB, SK, MSS, NPK, DV, FJV, HK, DO’H

Writing – original draft: YF, MY, IAW and BP

Writing – review & editing: All the authors reviewed and accepted the final contents of the manuscript.

## Competing interests

BP serves on the External Immunology Board of GSK and on the Scientific Advisory Board of Sanofi, Medicago, CircBio, Boehringer-Ingelheim, Icosavax, and EdJen. RSB serves on the Scientific Advisory Board of Takeda, VaxArt, and Invivyd and has collaborations with Janssen Pharmaceuticals, Gilead, Chimerix and Pardas Biosciences. NPK is a co-founder, shareholder, paid consultant, and chair of the scientific advisory board of Icosavax. The King laboratory has received unrelated sponsored research agreements from Pfizer and GSK. SS is an employee of the Sino Biological US company. DO’H is an employee of the GSK group of companies.

## Data and materials availability

All data associated with this study are present in the paper or the Supplementary Materials. AS03, RBD-NP, and Spike protein HexaPro are respectively available from GSK; Institute for Protein Design, University of Washington; and the University of Texas Austin under a material transfer agreement with the university or institution. X-ray coordinates and structure factors are currently being deposited in the RCSB Protein Data Bank and the identification numbers will be publicly available at the time of publication. Other materials are available from BP on request.

