## Supplementary materials for "Extremely potent pan-sarbecovirus neutralizing antibodies generated by immunization of macaques with an AS03-adjuvanted monovalent subunit vaccine against SARS-CoV-2"

**MATERIALS AND METHODS**

No statistical methods were used to predetermine the sample size. The experiments were not randomized. The investigators were not blinded to allocation during experiments and outcome assessment.

**Antigen-specific, memory B cell staining and single-cell sorting**

Banked PBMCs (*8, 9*) were thawed and washed twice with 10 mL of FACS buffer (1 x PBS containing 2% FBS and 1 mM EDTA) and resuspended in 100 μL of 1x PBS containing Zombie UV live/dead dye at 1:200 dilution (BioLegend, 423108) and incubated at room temperature for 15 minutes. Following washing, cells were incubated with an antibody cocktail for 1 hour protected from light on ice. The following antibodies were used: CD3 BV650 (BD Biosciences, 563916), CD14 BV650 (BioLegend, 301836), CD16 BV650 (BioLegend, 302042), CD20 APC-Cy7 (BioLegend, 302314), CD27 PE-Cy7 (BioLegend, 302838), IgM PerCP-Cy5.5 (BioLegend, 314512), IgD PE (Southern Biotech, 2030-09), IgG BUV395 (BD Biosciences, 564229) and AF488 labeled RBD of SARS-CoV-2 beta variant (SinoBiological, 40592-V08H85-B) and BV421 labeled spike (SinoBiological, 40589-V27B-B) or RBD (BioLegend, 793906) protein of SARS-CoV-2 Wuhan strain. All antibodies were used as per the manufacturer's instruction and the final concentration of each probe was 0.1 μg/ml. Single live CD3^-^ CD14^-^ CD16^-^ CD20^+^ IgM^-^ IgD^-^ IgG^+^ Probe^+^ B cells were sorted into individual wells of 96-well plates containing 16 μl of lysis buffer per well using a FACS Aria III and FACSDiva software (Becton Dickinson) for acquisition and FlowJo for analysis. The sorted cells were immediately frozen on dry ice and used for subsequent RNA reverse transcription as described below.

**Generation of recombinant humanized monoclonal antibodies**

Monoclonal antibodies were generated following established protocols (*53, 54*). In brief, single memory B cells were sorted with BD FACS Aria II into 96-well plates containing 16 μl of lysis buffer. The lysis buffer was composed of 20 U RNAse inhibitor (Invitrogen), 5 mM DTT (Invitrogen), 4 μl 5x RT buffer (Invitrogen), 0.0625 μl Igepal (Sigma), 10 μg/ml Carrier RNA (Applied Biosystems). The 96-well plates went through a quick freeze-thaw cycle, and 0.5 μg Oligo(dT)18 (Thermo Scientific), 0.5 mM dNTP mix (Invitrogen), and 200 U Superscript IV (Invitrogen) was added in a total volume of 4 μl followed by thorough mixing and spinning. The reverse transcription was performed as follows: 10 min at 42 ºC, 10 min at 23 ºC, 20 min at 50 ºC, 5 min at 55 ºC, 10 min at 80 ºC and finally cooling to 4 ºC. Ig heavy chain and light chain (kappa/lambda) rearrangements were amplified by nested PCR (HotStarTaq DNA Polymerase, QIAGEN) using primer cocktails (**Table S1**) specific for all V gene families and constant domains at a concentration of 250 nM per primer. The PCR mix consisted of 2.5 μl 10x PCR buffer, 0.5 μl 10 mM dNTP mix (Invitrogen), 0.5 μl 25 mM MgCl_2_ (only added in the first round PCR), 5 ul Q-solution, 1 U HotStarTaq, 0.5 ul 5’ and 3’ primers. Water was added up to a total volume of 25 μl. The PCR programs are shown in Table S2. The 2nd round PCR products were evaluated on 2% agarose gels, purified using QIAquick spin columns (Qiagen) and sequenced using 2nd round PCR reverse primers. The sequences were analyzed using the online IMGT/HighV-QUEST tool. The productive heavy-light paired Ig genes were used for antibody production (Sino Biological, High-throughput Antibody Production Service).

**Meso-scale electrochemiluminescence immunoassay (ECLIA)**

V-plex SARS-CoV-2 Panel 9 (human IgG) kit from Mesoscale Discovery (K15448U-2) was used to evaluate antibodies binding to RBD antigens from the following lineages: A (WT), (B.1.1.7), (B.1.214.2), (B.1.351), (B.1.427), (B.1.429), (B.1.525), (B.1.526), (B.1.526.1) (B.1.526.2), (B.1.617), (B.1.617.1), (B.1.617.3), (P.1), (P.3), and (R.1). The assay was performed as per the manufacturer’s instructions. Briefly, the multi-spot 96 well plates were blocked in 0.15 ml of blocking solution with shaking at 700 rpm at room temperature. After 30 min of blocking, 50 μl of monoclonal antibodies was added to each plate in the designated wells and incubated at room temperature for 2 h with shaking. Monoclonal antibodies were assayed at 10 μg/ml starting concentration and 11 additional 4-fold serial dilutions. Plates were washed and 50 μl of Sulfo-tag conjugated anti-IgG was added, and the plates were incubated at room temperature for 1 h. After incubation, the plates were washed, and 0.15 ml of MSD-Gold read buffer was added. The plates were immediately read using the MSD instrument (MESO QuickPlex SQ 120MM). The binding capacities of monoclonal antibodies were presented as the MSD arbitrary light unit area under curve (MSD AUC), calculated using Prism v9.3.1 (GraphPad).

**The enzyme-linked immunosorbent assay (ELISA)**

All monoclonal antibodies generated in this study were screened for their binding to the spikes of SARS-CoV-2 Wuhan, BA.1, and BA.4/5 strains by ELISA. Briefly, 96-well plates were coated with 50 μl per well of a 2 μg/ml spike protein solution in PBS overnight at 4 °C. Plates were washed 3 times with washing buffer (1× PBS with 0.05% Tween-20 (MP Biomedicals)) and incubated with 200 μl per well blocking buffer (1× PBS with 5% skim milk powder (Bio-Rad) and 0.05% Tween-20 (MP Biomedicals)) for 1 h at room temperature. Immediately after blocking, 100 μl of monoclonal antibodies was added to each plate in the designated wells and incubated at room temperature for 1 h. Monoclonal antibodies were assayed at 1 μg/ml starting concentration and 5 additional 5-fold serial dilutions. Plates were washed and 100 μl of horseradish peroxidase (HRP) conjugated goat anti-human IgG (Sigma, AP112P) in blocking buffer at a 1:5000 dilution was added, and the plates were incubated at room temperature for 1 h. After incubation, the plates were washed three times and 100 μl per well of TMB (Sigma, ES022-500ML) was added. The reaction was developed for 4 min and stopped by adding 100 μl per well of 450 nm stop solution (Abcam, ab171529). The plates were immediately read using an ELISA microplate reader (Bio-Rad). The binding capacities of monoclonal antibodies were presented as ELISA area under curve (AUC), calculated using Prism v9.3.1 (GraphPad software).

For experiments regarding the competition between mAbs for RBD binding, purified monoclonal antibodies (CR3022, CC12.3, CV07-270, S2X259, and S2M11) were biotinylated using Lightning-Link® Biotinylation Kit (Abcam, ab201795) according to the manufacturer’s instructions. 96-well plates were coated with 50 μl per well of a 2 μg/ml spike protein solution in PBS overnight at 4 °C. Plates were washed 3 times with washing buffer (1× PBS with 0.05% Tween-20 (MP Biomedicals)) and incubated with 200 μl per well blocking buffer (1× PBS with 5% skim milk powder (Bio-Rad) and 0.05% Tween-20 (MP Biomedicals)) for 1 h at room temperature Immediately after blocking, 100 μl of monoclonal antibodies was added to each plate in the designated wells and incubated at room temperature for 30 minutes. Monoclonal antibodies were assayed at 10 μg/ml starting concentration and 6 additional two-fold serial dilutions. Then, without washing, 100 μl of biotinylated monoclonal antibodies at 500 ng/ml was added and incubated for additional 30 minutes, followed by detection using HRP-conjugated streptavidin (Abcam, ab7403) and TMB (Sigma, ES022-500ML). Background by the HRP-conjugated detection antibodies alone was subtracted from all absorbance values.

**Pseudovirion neutralization assay (PsVNA)**

For initial screening, neutralization assays using pseudotyped viruses (using HIV backbone) carrying spikes of SARS-CoV-2 wild type, BA.1, and BA.4/5 strains, respectively, were performed by Sino Biological Inc. Briefly, 293T cells overexpressing ACE2 were seeded in 96-well plates (Costar, 3955) at 3 × 10^4^ cells/well. 50 μL of serially diluted mAbs or a control monoclonal antibody (S2H97) were mixed with 50 μL of pseudovirus and then they were added into plates seeded with 293T-ACE2 cells (1 × 10^4^ TCID50/mL). A positive control was set up using a mixture of 50 μL DMEM medium and 50 μL pseudovirus. 100 μL DMEM was used as a negative control. Cells were cultured at 37 °C for 64 h. Luminous value was detected by Luminometer (Berthold Technologies, Centro LB 960) and the inhibitory rate was calculated by (1 − (mean RLU of sample − RLU of negative control) / (RLU of positive control − RLU of negative control) *100%). Experiments were performed in duplicate.

The 15 most potent BA.1 neutralizing antibodies were evaluated in a qualified SARS-CoV-2 PsVNA performed in the laboratory of U.S. Food and Drug Administration. Pseudovirions were produced as previously described (*55*). Briefly, human codon-optimized cDNA encoding SARS-CoV-2 spike glycoprotein of the WA-1/2020 and variants and SARS-CoV spike glycoprotein were synthesized by GenScript and cloned into eukaryotic cell expression vector pcDNA 3.1 between the BamHI and XhoI sites. Pseudovirions were produced by co-transfection Lenti‐X 293T cells with psPAX2(gag/pol), pTrip-luc lentiviral vector and pcDNA 3.1 SARS-CoV-2-spike-deltaC19, using Lipofectamine 3000. The supernatants were harvested at 48h post-transfection, filtered through 0.45µm membranes, and titrated using 293T-ACE2-TMPRSS2 cells (HEK 293T cells expressing ACE2 and TMPRSS2 proteins).

Neutralization assays were performed as previously described (*56, 57*). For the neutralization assay, 50 µL of pseudovirions (counting ~200,000 relative light units) were pre-incubated with an equal volume of medium containing serial dilutions of mAbs at room temperature for 1h. Then 50 µL of virus and antibody or serum mixtures were added to 293T-ACE2-TMPRSS2 cells (10^4^ cells/50 μL) in a 96-well plate. Controls included cells-only control, virus without any antibody control. After a 3 h incubation, fresh medium was added to the wells. Cells were lysed 24 h later, and luciferase activity was measured using One-Glo luciferase assay system (Promega, Cat# E6130). Each assay was performed in duplicate, and the 50% neutralization titer was calculated using Prism 9 (GraphPad Software).

**Production of Viruses for in vitro neutralization and in vivo challenge assays**

All recombinant infectious clone-based viruses were approved by the University of North Carolina at Chapel Hill Institutional Review Board under Schedule G 78684, 100475, 60350, 104615, and 11515. Viruses were designed using our previously described infectious clone systems to incorporate mutations that were previously defined as necessary for effective pathogenic infection of mice. These mouse-adapted mutations (MA) were used for the design of the SARS-CoV-2 in vivo challenge viruses (MA10), as well as for our SARS-CoV and SHC014 (MA15) viruses. Additionally, viruses used in live virus neutralization assays were derived from infectious clones where the ORF7 gene was replaced with a nano-luciferase gene cassette.

Viruses were derived following the ligation of cDNA of infections clone fragments, followed by in vitro transcription with mMessage Machine T7 polymerase (ThermoFisher). A separate reaction using a T7 promoter upstream of the nucleocapsid gene was used to produce nucleocapsid mRNA to aid in virus replication and recovery. Prior to electroporation, cells were mixed with the full-length mRNA as well as the nucleocapsid mRNA. Vero E6 cells over-expressing human TMPRSS2/ACE2 cells in phosphate buffered saline (PBS, Gibco) were electroporated under the following conditions: 450 volts, 50 microfarads, 4 pulses and allowed to recover for 10 minutes. The cells were then plated into a T75 flask. Passage 0 (p0) stocks were recovered ~24-36 hours post-electroporation. Passage 1 working stocks were created by inoculation of a confluent T175 flask of Vero E6 TMPRSS2/ACE2 cells with 1 mL of p0 virus. Stocks were then titered via plaque assay where virus was serially diluted ten-fold and inoculated onto confluent monolayers of Vero E6 cells in 6 well plates. Plates were incubated for 1 hour with gentle rocking every 15 minutes. Subsequently, Dulbecco’s modified Eagle’s medium (DMEM; Gibco) with Fetal Clone II serum (FCII, Hyclone), and 1X antibiotic/antimycotic (Gibco), and 0.8% agarose was applied as an overlay.

**Live-virus neutralization assays**

Antibodies were diluted 1:20 by adding 11.25 μL of antibodies to 213.75 μL DMEM supplemented with 10% Fetal Clone II. Antibodies were then serially diluted 1:3 seven times by adding 75 μL of the previous dilution into 150 μL of media with 75 μL removed from the final well. Next, 150 μL of media containing 1600 PFU/mL of viruses expressing nanoluciferase (nLuc) was mixed with the diluted antibodies and allowed to incubate for 1 hr at 37℃, after which 100 μL of the virus + antibody mix was added to individual wells in a 96 well plate seeded 24 hours prior with 2x10^4^ cells for a final 800 PFU virus per well. Plates were incubated for 48 hours at 37℃ with 5% CO2. After incubation, luciferase activity was measured with the Nano-Glo Luciferase Assay System (Promega) according to the manufacturer specifications. Neutralization titers (IC_50_) were defined as the dilution at which a 50% reduction in RLU was observed relative to the virus (no antibody) control.

**Mouse & In vivo challenges**

All animal work was approved by Institutional Animal Care and Use Committee at University of North Carolina at Chapel Hill under protocol 20-114 and 20-200 according to guidelines outlined by the Association for the Assessment and Accreditation of Laboratory Animal Care and the U.S. Department of Agriculture. All virus studies were performed in animal biosafety level 3 facilities at University of North Carolina at Chapel Hill.

Female BALB/c mice were obtained from Envigo (strain 047). Twelve hours prior to infection mice received 200 μg of indicated mAbs or an isotype recombinant human anti-Dengue envelope antibody (2D22) intraperitoneally. The following morning, mice were infected intranasally under ketamine/xylazine anesthesia with plaque-forming unit doses of 1x10^3^ (SARS-CoV-2 MA10), 1x10^4^ (SARS-CoV MA15), or 1x10^5^ (SARS-CoV-2 BA.1 and SARS-CoV MA15 SHC014) in 50 μL of PBS. At indicated timepoints, a subset of mice were euthanized by isoflurane overdose, and lung tissue was harvested for titer, gross lung discoloration, and histopathological analyses. Titer and samples were stored at −80°C, and histopathology samples at 4°C in 10% phosphate buffered saline. Gross lung discoloration scores were based on the number and severity of visible lung surface hemorrhage at the time of harvest (e.g., dark red vs. anatomical pink coloring).

**Biolayer Interferometry Binding Assay**

RBD proteins for the biolayer interferometry (BLI) binding assay were expressed in human cells. RBDs were cloned into phCMV3 vector and fused with a C-terminal His_6_ tag. The plasmids were transiently transfected into Expi293F cells using ExpiFectamine 293 reagent (Thermo Fisher Scientific) according to the manufacturer’s instructions. The supernatant was collected at 7 d post-transfection. The His_6_-tagged proteins were then purified with Ni Sepharose Excel protein purification resin (Cytiva) followed by size exclusion chromatography. Omicron RBD was purchased from ACROBiosystems Inc.

The BLI assays were performed using an Octet Red instrument (FortéBio) as described previously (*26*). To measure the binding kinetics of mAbs and RBDs, the mAbs were diluted with kinetic buffer (1× phosphate-buffered saline [pH 7.4], 0.01% bovine serum albumin and 0.002% Tween 20) into 15 µg/mL. The mAbs were then loaded onto anti-human IgG Fc (AHC) biosensors and interacted with 100 nM of RBDs. The assay consisted of the following steps: 1) baseline, 1 min with 1× kinetic buffer; 2) loading, 90 s with mAbs; 3) wash, 15 s wash of unbound mAbs with 1× kinetic buffer; 4) baseline, 1 min with 1× kinetic buffer; 5) association, 90 s with RBDs; and 6) dissociation, 90 s with 1× kinetic buffer. For estimating the dissociation constant (KD), a 1:1 binding model was used.

**Crystallization and structural determination**

Expression and purification of the SARS-CoV-2 spike receptor-binding domains (RBDs) for crystallization were as described previously (*18*). Briefly, wild-type and BA.2 SARS-CoV-2 RBDs (residues 333-529) of the spike (S) proteins were cloned into a customized pFastBac vector (*58*) and fused with an N-terminal gp67 signal peptide and C-terminal His_6_ tag. The recombinant bacmid DNAs were generated using the Bac-to-Bac system (Life Technologies). Baculoviruses were generated by transfecting purified bacmid DNAs into Sf9 cells using FuGENE HD (Promega), and subsequently used to infect suspension cultures of High Five cells (Life Technologies) at an MOI of 5 to 10. Infected High Five cells were incubated at 28 °C with shaking at 110 rpm for 72 h for protein expression. The supernatants were then concentrated using a 10 kDa MW cutoff Centramate cassette (Pall Corporation). The RBD proteins were purified by Ni-NTA, followed by size exclusion chromatography, and buffer was exchanged into 20 mM Tris-HCl pH 7.4 and 150 mM NaCl.

25F9/RBD (WT), 21B6/RBD (WT), 20A7/RBD (WT), and 20A7/RBD (BA.2) complexes were formed by mixing each of the protein components in an equimolar ratio and incubating overnight at 4°C. The protein complexes were adjusted to 11–12 mg/ml and screened for crystallization using the 384 conditions of the JCSG Core Suite (Qiagen) on our robotic CrystalMation system (Rigaku) at Scripps Research. Crystallization trials were set-up by the vapor diffusion method in sitting drops containing 0.1 μl of protein and 0.1 μl of reservoir solution. For the 25F9/RBD (WT) complex, optimized crystals were grown in drops containing 1.6 M ammonium sulfate, 0.1 M bicine pH 9, and 15% glycerol at 20°C. Crystals appeared on day 14 and were harvested on day 28 by soaking in reservoir solution supplemented with 15% (v/v) glycerol. Diffraction data were collected at cryogenic temperature (100 K) at beamline 23-ID-D of the Advanced Photon Source (APS) at Argonne National Labs. For the 21B6/RBD (WT) complex, optimized crystals were grown in drops containing 0.1 M sodium citrate, pH 3.3 and 1.45 M ammonium sulfate at 20°C. Crystals appeared on day 7 and were harvested on day 15 by soaking in reservoir solution supplemented with 20% (v/v) ethylene glycol. Diffraction data were collected at cryogenic temperature (100 K) at beamline 23-ID-B of the APS at Argonne National Labs. For the 20A7/RBD (WT) complex, optimized crystals were grown in drops containing 0.2 M CaCl_2_, 10% ethylene glycol (v/v), and 20% polyethylene glycol 3350 (w/v) at 20°C. Crystals appeared on day 7 and were harvested on day 15 with no additional cryoprotectant. Diffraction data were collected at cryogenic temperature (100 K) at beamline 23-ID-B of the APS at Argonne National Labs. For the 20A7/RBD (BA.2) complex, optimized crystals were then grown in drops containing 0.1 M HEPES pH 7.5, 10% (v/v) glycerol, 5% (w/v) polyethylene glycol 3000, and 30% (v/v) polyethylene glycol 400 at 20°C. Crystals appeared on day 7 and were harvested on day 10 by soaking in reservoir solution supplemented with 15% (v/v) ethylene glycol. Diffraction data were collected at cryogenic temperature (100 K) at the Stanford Synchrotron Radiation Lightsource (SSRL) on Scripps/Stanford beamline 12-1. Diffraction data were processed with HKL2000 (*59*). Structures were solved by molecular replacement using PHASER (*60*). Iterative model building and refinement were carried out in COOT (*61*) and PHENIX (*62*), respectively. Epitope and paratope residues, as well as their interactions, were identified by accessing PISA at the European Bioinformatics Institute (http://www.ebi.ac.uk/pdbe/prot_int/pistart.html) (*63*).

**Statistical analysis**

Individual-level data for all the figures are presented in Data file S1. The difference between any two groups at a time point was measured using a two-tailed nonparametric Mann-Whitney unpaired rank-sum test or the two-tailed Kruskal–Wallis test with subsequent Dunn’s multiple-comparisons test. The difference between groups at different time points was measured using two-way ANOVA. The difference between multiple time points was measured using one-way ANOVA. The difference between different categories was measured by the two-tailed chi-square test. All correlations were Spearman’s correlations based on ranks. All statistical analyses were performed using GraphPad Prism v.9.0.0 or R version 3.6.1. All the figures were made in GraphPad Prism or R and organized in Adobe Illustrator.

**Supplementary Figures**

**Figure S1. Comparison of memory B cell responses in different adjuvant groups.**

(**A**) Schematic representation of the study design. (**B**) Gating strategy of analyzing the percentage of spike^+^ and/or RBD^+^ B cells in total CD20^+^ cells. Gating was on singlets that were live CD20^+^ CD3^-^ CD14^-^ CD16^-^ IgM^-^/D^-^ IgG^+^ Antigen^+^. (**C**) Dot plot summarizes the percentage of RBD-binding B cells in total CD20^+^ B cells in 21 vaccinated individuals. Horizontal bars indicate mean values. Statistical significance between groups was determined using two-tailed Mann–Whitney U tests. n = 3 (Alum), 5 (AS03 and AS37), and 4 (CpG-Alum and SWE). (**D**) Graphs show the somatic hypermutation rates of the productive IGHV (left) and IGLV (right) genes isolated from animals vaccinated with RBD-NP plus different adjuvants at day 42. The boxes inside the violin plot show median, upper, and lower quartiles. Each dot represents an individual gene. For heavy chain, n = 22 (Alum), 42 (OW), 44 (AS03), 74 (AS37), 29 (CpG) and for light chain, n = 14 (Alum), 35 (OW), 37 (AS03), 78 (AS37), 40 (CpG). (**E**) as in (D), but for CDR3 length in amino acids (aa). The boxes inside the violin plot show median, upper, and lower quartiles. Each dot represents an individual gene. For heavy chain, n = 22 (Alum), 42 (OW), 44 (AS03), 74 (AS37), 29 (CpG) and for light chain, n = 14 (Alum), 35 (OW), 37 (AS03), 78 (AS37), 40 (CpG).

**Figure S2. Antibody and memory B cell response.**

(**A**) Frequency of antigen (spike- or RBD-) specific memory B cells relative to CD20^+^ B cells is shown for samples from RBD-NP (blue) or Hexapro-NP (red) group. Paired samples are connected with a dashed line. Geometric means are presented in thick lines, while shades are for a 95% confidence interval. (**B**) The graph shows the relative abundance of IGVH gene usage in antigen-specific memory B cells in Rhesus macaques calculated using the IMGT database. (**C**) The graph shows the relative abundance of IGVH gene usage in antigen-specific memory B cells in Rhesus macaques calculated using the KIMDB database. (**D**) IgBlast analysis of human VH3-53 germline gene and human VH4-59 germline gene using the IGMT rhesus macaque database.

**Figure S3. Binding profiles of monoclonal antibodies.**

(**A**) Graphs show results of Meso scale discovery (MSD) ECLIAs measuring cross-reactive binding profiles of monoclonal antibodies isolated from individuals vaccinated with RBD-NP plus different adjuvants. Data were presented in area under curve (AUC). Black horizontal bars represent the median. Statistical significance was determined using Tukey’s multiple comparisons test. P values were labeled on top. (**B**) Graphs show anti-SARS-CoV-2 Wuhan spike ELISA titration curves for all 514 monoclonal antibodies generated in this study. Monoclonal antibodies showing OD450 above 1.0 at 1 μg/ml concentration were highlighted in red. The Spearman’s correlation between the binding AUC and somatic hypermutation rates (SHM) of IGHV genes of the highlighted antibodies was shown on the right. (**C**) Graphs show the anti-SARS-CoV-2 Wuhan spike activity (AUC) of the highlighted monoclonal antibodies in (B) at indicated time points. The boxes show the median, upper, and lower quartiles. The whiskers show min to max. Each dot represents one antibody. The statistical differences between time points were calculated using one-way ANOVA. P values were shown on top.

**Figure S4. Cross-reactive properties of monoclonal antibodies.**

(**A**) Heatmaps show the binding activities of the red highlighted antibodies in Fig. S3B against spikes of SARS-CoV-2 Wuhan, BA.1, and BA.4/5. Antibodies isolated at indicated time points were grouped (the number of antibodies per group was shown in brackets). The gradient color bar indicated the BA.1 binding capacity (area under curve, AUC) from 40 to 4000. (**B**) Graphs show anti-SARS-CoV-2 BA.1 spike ELISA titration curves for all 514 monoclonal antibodies generated in this study. Monoclonal antibodies showing OD450 above 1.0 at 1 μg/ml concentration were highlighted in red. The Spearman’s correlation between the BA.1 binding AUC and somatic hypermutation rates (SHM) of IGHV genes of the highlighted antibodies was shown on the right. (**C**) Graphs show the anti-SARS-CoV-2 BA.1 spike activity (AUC) of the highlighted monoclonal antibodies in (B) at indicated time points. The boxes show the median, upper, and lower quartiles. The whiskers show min to max. Each dot represents an antibody. The statistical differences between time points were calculated using one-way ANOVA. P values were shown on top. (**D**) Graphs show anti-SARS-CoV-2 BA.4/5 spike ELISA titration curves for all 514 monoclonal antibodies generated in this study. Monoclonal antibodies showing OD450 above 1.0 at 1 μg/ml concentration were highlighted in red. The Spearman’s correlation between the BA.4/5 binding AUC and somatic hypermutation rates (SHM) of IGHV genes of the highlighted antibodies was shown on the right. (**E**) Graphs show the anti-SARS-CoV-2 BA.4/5 spike activity (AUC) of the highlighted monoclonal antibodies in (D) at indicated time points. The boxes show the median, upper, and lower quartiles. The whiskers show min to max. Each dot represents an antibody. The statistical differences between time points were calculated using one-way ANOVA. P values were shown on top. (**F**) The antibodies were categorized based on the cross-reactive properties as shown in the right legends. Pie charts illustrate the fraction of antibodies in each category at indicated time points and the number of antibodies per group was the same as in (A).

**Figure S5. Binding and neutralizing activities of the top 15 SARS-CoV-2 BA.1 neutralizing monoclonal antibodies.**

(**A**) Monoclonal antibodies showing comparable or better SARS-CoV-2 BA.1 neutralizing potency (lower half maximal inhibition concentration, IC50, ug/ml) than that of a control antibody, S2H97 (purple) were highlighted in red and selected for further characterization. The numbers within the graphs show GMTs. The statistical differences between viruses were calculated using one-way ANOVA test (*P < 0.05, ****P < 0.0001). (**B**) Neutralization of 15 mAbs against pseudotyped SARS-CoV-2 (WA1), 4 previous SARS-CoV-2 variants of concern [B.1.1.7 (Alpha), B.1.351 (Beta), P.1 (Gamma), B.1.617.2 (Delta)], current variants of concern SARS-CoV-2 Omicron sublineage (BA.1, BA.2, BA.3, BA.4/5) and SARS-CoV. Data are presented as the mean ± SD. (**C**) Percent neutralization of mAbs against authentic SARS-CoV-2 D614G, Omicron BA.1, and Pangolin, SARS-CoV, WIV1, SHC014, and MERS-CoV was presented as IC50 (ng/ml). (**D**) Pseudovirus neutralization IC50 values for mAbs against BQ.1, BQ.1.1 and XBB subvariants.

**Figure S6. Binding Epitope Characterization of the 7 selected ultrapotent monoclonal antibodies.**

(**A**) Competition ELISA (blockade-of-binding) between selected antibodies (labeled in different shapes and colors) at serially diluted concentration and the indicated biotinylated antibodies (shown as the title) with known binding epitopes at a fixed concentration of 0.5 μg/ml. Controls (green) show the absorbance values with only the detection antibody. (**B**) Heatmap illustrates the competition for SARS-CoV-2 spike binding between combinations of selected monoclonal antibodies. Shades and percentages in squares indicate the degree of competition for spike binding of detection antibodies (row) in the presence of mAbs (column) at a concentration of 10 μg/ml.

**Figure S7. Crystal structure of 21B6 in complex with SARS-CoV-2 RBD.**

The heavy and light chains of 21B6 variable domains are in dark and light teal. (**A**) Crystal structure of 21B6 in complex with SARS-CoV-2 RBD. (**B**) The surface area of SARS-CoV-2 buried by heavy and light chains of 21B6. (**C**) 21B6 interacts with RBD with CDRs H1, H2, H3, L2, and HFR1. (**D**) SARS-CoV-2 RBD in complex with 21B6 superimposed onto an RBD/hACE2 complex structure (PDB 6M0J) shows that 21B6 would clash (red circle) with the hACE2 receptor. (**E**) Sequence alignment of epitope residues (defined as BSA > 0 Å^2^) in a subset of SARS-like viruses. Residues that differ from wild-type SARS-CoV-2 are indicated in red. (F-I) Molecular interactions between RBD and 21B6. Hydrogen bonds and salt bridges are indicated by dashed lines.

**Figure S8. Structural convergence of rhesus macaque IGHV3-73-encoded antibodies targeting SARS-CoV-2 RBD.**

The SARS-CoV-2 RBDs are shown in whitish grey, while the antibody heavy and light chains are in orange and yellow. Hydrogen bonds and salt bridges are indicated by dashed lines. (**A**) Rhesus macaque IGHV3-73-encoded antibody 20A7 adopts the same binding mode as two previously published antibodies (**B**) K288.2 and (**C**) K398.22 encoded by the same VH germline gene (*22*). (**D-F**) Residue V_H_ E33 of all three IGHV3-73-encoded antibodies (D) 20A7, (E) K288.2, and (F) K398.22 hydrogen bond with the backbone amide of RBD-V503. (**G**) Sequence alignment of all IGHV3-73 alleles available in the macaque KIMDB Ig database. Residues at position 33 are highlighted in yellow. (**H**) BLI assay of 20A7 wild-type Fab and an E33V mutant binding to SARS-CoV-2 RBD.

**Figure S9. Epitope map of SARS-CoV-2 RBD.**

The receptor binding site (RBS) and CR3022 epitope are shown in green and pink, respectively. RBS-D is located on one corner of the RBS and is indicated by a black circle. The epitopes of 20A7, 25F9, and SA55 are outlined in orange, blue, and purple, respectively. Epitope residues are defined as BSA > 0 Å^2^ and calculated from RBD structures with CR3022 (PDB 6W41), ACE2 (PDB 6M0J), SA55 (PDB 7Y0W) and 20A7 and 25F9 from this study.

**Supplementary Tables**

**Table S1. Primer list**

| Primer | 5’ – 3’ Sequence | Source |
| --- | --- | --- |
| 5′VH1.L1 | ATGGACTKGACCTGGAGG | Sundling 2012 (*50*) |
| 5'VH1/7 | GGACCTGACCCGGAGGATC | modified Wiehe 2014 (*51*) |
| 5′VH2.L1 | ATGGACACGCTTTGCTCC | Sundling 2012 |
| 5′VH3A.L1 | ATGGAGTTKGGGCTGAGCTG | Sundling 2012 |
| 5'VH3_EXT2 | GGGGCTGAGYTGGGTTTTC | modified Wiehe 2014 |
| 5'VH3_EXT4 | TGGGCTGAGCTKGGTTTTY | modified Wiehe 2014 |
| 5′VH3B.L1 | ATGGAGTTTGKRCTGAGCTGG | Sundling 2012 |
| 5′VH3C.L1 | ATGGAGTCRTGGCTGAGCTG | modified Sundling 2012 |
| 5′VH3D.L1 | ATGGAGTTTGTGCTGAGTTTGG | Sundling 2012 |
| 5'VH4_EXT1 | ATGAAGCACCTGTGGTTCTBC | modified Wiehe 2014 |
| 5'VH4_EXT2 | ATGAAGCACCTGKGGTTCTTY | modified Wiehe 2014 |
| 5'VH5_EXT | ATGGGGTCAACTGCCMTCC | modified Wiehe 2014 |
| 5'VH6_EXT | ATGTCTGTCTCCTTCCTCATCGTC | modified Wiehe 2014 |
| 3' IgG_ext | TGTGCACGCCGCTGGTCAG | modified Wiehe 2014 |
| 5' VK1A_ext | TGTGACATCCAGATGACCCAG | New design |
| 5' VK1B_ext | GTGCCAGATGTGACATTCAG | New design |
| 5' VK2A_ext | ATGAGGCTCCCTGCTCAGCTC | New design |
| 5' VK2B_ext | CTSCCTGCTCWGCTCCTG | New design |
| 5' VK2C_ext | GATCCASTGGGGATGTTGY | New design |
| 5' VK3A_ext | CAGCACAGCTTCTCTTCCTCCTG | New design |
| 5' VK3B_ext | AGCTCGGCTTCTCTGCCTTCT | New design |
| 5' VK3C_ext | AGCTCAGCTTCTCTTCCTCCTGC | New design |
| 5' VK4A_ext | ATGGTGTCACAGACCCAAGTCTT | New design |
| 5' VK4B_ext | ATGGTGCTACAGACCCAGGTCCT | New design |
| 5' VK5_ext | GGTTCASCTCCTCAGCTTCCTC | New design |
| 5' VK6_ext | TTCTGCTSCTCTGGGTTCCAG | New design |
| 5' VK7_ext | ATGGGGTCCTGGGCTCCTT | New design |
| 3' CK_ext | GTCCTGCTCTGTGACACTCTCCT | modified Sundling 2012 |
| 5' VL1A-ext | ATGGCCTGGTYYCCTCTC | Sundling 2012 |
| 5' VL1B_ext | GGTCCTGGGCCCAGTCTG | New design |
| 5' VL2/7/10_ext | ATGSYCTGGRCTCTGCTCCTC | New design |
| 5' VL3A_ext | ACAGGTTCYGTGGTTTCYTCTG | New design |
| 5' VL3B_ext | ATGGCCTGGATTCCTCTCCT | modified Sundling 2012 |
| 5' VL3C_ext | ATGGCCTGGACCCYTCTCCT | modified Wiehe 2014 |
| 5' VL4A_ext | ATGGCCTGGGTCTCCTTC | Sundling 2012 |
| 5' VL4B_ext | ATGGCCTGGACCCCACTC | Sundling 2012 |
| 5' VL5/11_ext | ATGGCCTGGACHCCTCTCCT | modified Sundling 2012 |
| 5' VL6_ext | ATGGCCTGGGCTCCACTCC | Sundling 2012 |
| 5' VL8_ext | ATGGCCTGGATGATGCTTCT | modified Sundling 2012 |
| 5' VL9_ext | ATGGCCTGGGCTCCTCTGCT | modified Sundling 2012 |
| 3' CL_ext | TGTTGTTGCTCTGTTTGGAGGG | Zhang 2019 (64) |
| 5'VH1a_int | CAG GTS CAG CTG GTG CAR TC | New design |
| 5'VH1b_int | GAG GTC CAG CTG GTG CAG TC | New design |
| 5'VH1c_int | CAG CTG GTG CAA TCC GGG | New design |
| 5'VH2_int | CAG GTS ACC TTG AAG GAG TC | New design |
| 5VH3a_int | GAS GTG CAG CTG GTR GAG TC | New design |
| 5'VH3b_int | GAG GTG CAG CTG GTG GMG TM | New design |
| 5'VH3c_int | GAR GTG CAG TTG GTG GAG TC | New design |
| 5'VH4a_int | CAG STG CAG CTG CAG GAG TC | New design |
| 5'VH4b_int | CAG GTG AAG CTG CAG CAG TG | New design |
| 5'VH5/7a_int | SAG GTG CAG CTG GTG CAG TC | New design |
| 5'VH6_int | CAG GTG CAG CTG CAG GAG TC | New design |
| 3' IgG_int | GAA GTA GTC CTT GAC CAG GCA | New design |
| 5' VK1a_int | GACATCCAGATGWCCCAGKCT | New design |
| 5' VK1b_int | GACATTCAGWTGWCCCAGTCTC | New design |
| 5' VK2a_int | GATATTGTGATGAYCCAGACTCC | New design |
| 5' VK2b_int | TGGGGATGTTGTGATGACTCAG | New design |
| 5' VK3a_int | CCTGCTGCTCTGGMTCCCA | New design |
| 5' VK3b_int | CCTGCTACTSTGGCTCCCAG | New design |
| 5' VK3c_int | CCTGCTACCTTGGCTCCCAG | New design |
| 5' VK4_int | GACATTGTGATGACCCAGTCTCC | Zhang 2019 |
| 5' VK5_int | TCTCTGATGCCAGGGCAGAAA | New design |
| 5' VK6_int | CTCTGGGTTCCAGYCTCCA | New design |
| 5' VK7_int | GACATTGTGCTGACCCAGTCTC | Zhang 2019 |
| 3' CK_int | ATTCAGCAGGCACACAACAGAG | Zhang 2019 |
| 5' VL1a/10_int | CAGGCAGGGCTGACTCAG | modified Zhang 2019 |
| 5' VL1b_int | CAGTCTGTGCTGACDCAGC | modified Zhang 2019 |
| 5' VL2_int | TCCTGGGCTCAGKCTGCC | New design |
| 5' VL3a_int | TCYTCTGRGCTGACTCAGGA | New design |
| 5' VL3b_int | TCCTMTGAKCTGACTCAGCCAC | New design |
| 5' VL4/5/9/11_int | CWGCCTGTGCTGACTCAGYC | New design |
| 5' VL6_int | GAGGTTGTGTTCACTCAGCCC | modified Zhang 2019 |
| 5' VL7_int | CAGGCTGTAGTGACTCAGGAGCC | modified Zhang 2019 |
| 5' VL8_int | GAGACTGTGGTGACCCAGGAGC | modified Zhang 2019 |
| 3' CL_int | CTCCCGGGTAGAAGTCACTGATC | New design |

**Table S2 PCR programs**

a. First round PCR program

| **Temperature** | **Duration** | **Number of Cycles** |
| --- | --- | --- |
| 95 ℃ | 5 min |  |
| 94-67-72 ℃ | 30-45-60 s | 3 |
| 94-64-72 ℃ | 30-45-60 s | 3 |
| 94-61-72 ℃ | 30-45-60 s | 3 |
| 94-58-72 ℃ | 30-45-60 s | 3 |
| 94-55-72 ℃ | 30-45-60 s | 3 |
| 94-52-72 ℃ | 30-45-60 s | 25 |
| 72 ℃ | 7 min |  |
| 4 ℃ | ∞ |  |

b. Second round PCR program

| **Temperature** | **Duration** | **Number of Cycles** |
| --- | --- | --- |
| 95 ℃ | 5 min |  |
| 94-58-72 ℃ | 30-45-60 s | 3 |
| 94-55-72 ℃ | 30-45-60 s | 3 |
| 94-52-72 ℃ | 30-45-60 s | 40 |
| 72 ℃ | 7 min |  |
| 4 ℃ | ∞ |  |

**Table S3. X-ray data collection and refinement statistics**

| **Data collection** | 25F9 + wild-type  SARS-CoV-2 RBD | 21B6 + wild-type  SARS-CoV-2 RBD | 20A7 + wild-type  SARS-CoV-2 RBD | 20A7 + BA.2  SARS-CoV-2 RBD |
| --- | --- | --- | --- | --- |
| Beamline | APS23ID-D | APS23ID-B | APS23ID-B | SSRL12-1 |
| Wavelength (Å) | 1.0332 | 1.0332 | 1.0332 | 0.9795 |
| Space group | P 1 | P 2_1_ 2_1_ 2_1_ | P 2_1_ 2_1_ 2_1_ | P 2_1_ 2_1_ 2_1_ |
| Unit cell parameters |  |  |  |  |
| a, b, c (Å) | 92.5, 105.8, 118.2 | 72.7, 78.8, 131.8 | 44.8, 131.4, 174.1 | 44.5, 132.2, 170.5 |
| α, β, γ (°) | 83.1, 67.5, 64.0 | 90, 90, 90 | 90, 90, 90 | 90, 90, 90 |
| Resolution (Å) ^a^ | 50.0-3.05 (3.10-3.05) | 50.0-1.75 (1.78-1.75) | 50.0-2.58 (2.62-2.58) | 50.0-2.30 (2.34-2.30) |
| Unique reflections ^a^ | 69,365 (3,475) | 75,221 (7,384) | 33,431 (3,097) | 45,705 (4,413) |
| Redundancy ^a^ | 2.7 (2.6) | 11.7 (9.2) | 11.6 (7.0) | 11.8 (7.7) |
| Completeness (%) ^a^ | 98.6 (97.5) | 97.7 (96.7) | 99.2 (92.0) | 99.7 (97.3) |
| <I/σ_I_> ^a^ | 1.8 (0.4) | 20.9 (1.1) | 13.4 (1.0) | 20.9 (1.2) |
| *R*_sym_^b^ (%) ^a^ | 43.5 (>100) | 12.8 (93.5) | 17.0 (82.6) | 10.9 (>100) |
| *R*_pim_^b^ (%) ^a^ | 30.9 (>100) | 3.9 (30.3) | 5.1 (31.1) | 4.3 (42.4) |
| CC_1/2_^c^ (%) ^a^ | 87.7 (37.9) | 99.8 (80.5) | 99.1 (68.2) | 99.6 (75.6) |
| **Refinement statistics** | | |  |  |
| Resolution (Å) | 46.2-3.35 | 39.4-1.75 | 41.3-2.58 | 43.1-2.30 |
| Reflections (work) | 52,773 | 75,149 | 33,383 | 45,687 |
| Reflections (test) | 2,519 | 1,993 | 1,564 | 2,314 |
| *R*_cryst_^d^ / *R*_free_^e^ (%) | 27.2/31.9 | 19.4/22.1 | 22.4/24.8 | 19.6/22.6 |
| No. of antibody/RBD copies in ASU | 4 | 1 | 1 | 1 |
| No. of atoms | 19,634 | 5,459 | 4,911 | 5044 |
| RBD | 6,302 | 1,557 | 1,503 | 1,569 |
| Fab | 13,253 | 3,309 | 3,233 | 3,277 |
| Ligands ^f^ | 79 | 10 | 0 | 25 |
| Solvent | 0 | 583 | 175 | 194 |
| Average *B-*values (Å^2^) | 73 | 31 | 55 | 56 |
| RBD | 76 | 32 | 71 | 78 |
| Fab | 72 | 29 | 48 | 46 |
| Ligands ^f^ | 70 | 41 | N/A | 56 |
| Solvent | N/A | 40 | 50 | 52 |
| Wilson *B*-value (Å^2^) | 39 | 24 | 51 | 45 |
| **RMSD from ideal geometry** | | |  |  |
| Bond length (Å) | 0.012 | 0.007 | 0.002 | 0.002 |
| Bond angle (^o^) | 0.64 | 1.1 | 0.54 | 0.55 |
| **Ramachandran statistics (%) ^g^** | | |  |  |
| Favored | 95.6 | 96.2 | 96.2 | 97.6 |
| Outliers | 0.20 | 0.32 | 0.17 | 0.00 |

^a^ Numbers in parentheses refer to the highest resolution shell.

^b^ *R*_sym_ = Σ*_hkl_* Σ*_i_* | I*_hkl,i_* - <I*_hkl_*> | / Σ*_hkl_* Σ*_i_* I*_hkl,i_* and R*_pim_* = Σ*_hkl_* (1/(n-1))^1/2^ Σ*_i_* | I*_hkl,i_* - <I*_hkl_*> | / Σ*_hkl_* Σ*_i_* I*_hkl,i_*, where I*_hkl,i_* is the scaled intensity of the i^th^ measurement of reflection h, k, l, <I*_hkl_*> is the average intensity for that reflection, and *n* is the redundancy.

^c^ CC_1/2_ = Pearson correlation coefficient between two random half datasets.

*^d^ R*_cryst_ = Σ*_hkl_* | *F*_o_ - *F*_c_ | / Σ*_hkl_* | *F*_o_ | x 100, where *F*_o_ and *F*_c_ are the observed and calculated structure factors, respectively.

^e^ *R*_free_ was calculated as for *R*_cryst_, but on a test set comprising 2.5% or 5% of the data excluded from refinement.

^f^ Bound ligands are SO_4_, ethylene glycol, glycerol, bicine, and PEG.

^g^ From MolProbity (*65*).
