## Supplementary Figures for "Extremely potent pan-sarbecovirus neutralizing antibodies generated by immunization of macaques with an AS03-adjuvanted monovalent subunit vaccine against SARS-CoV-2"

### Supplementary Figure 1

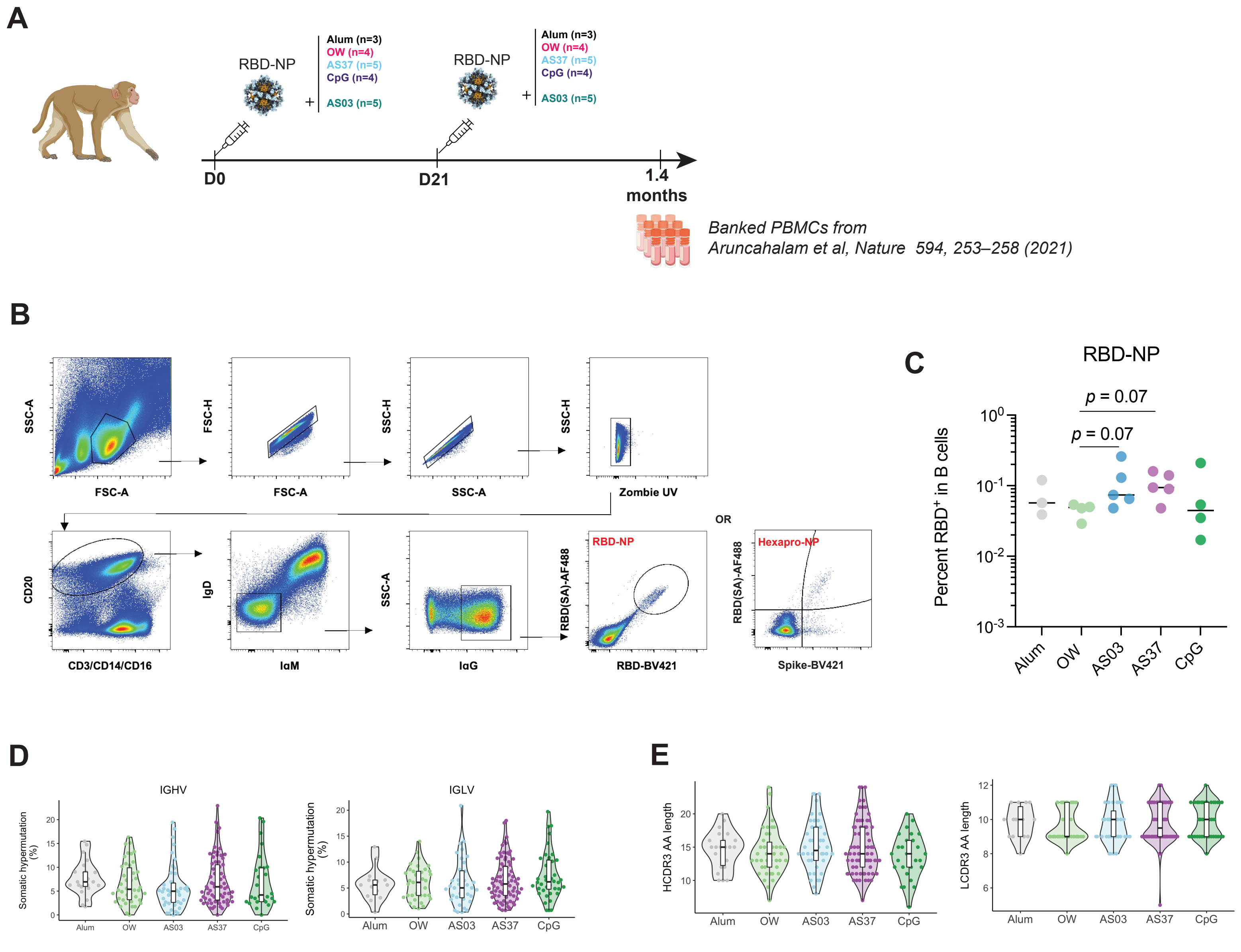

### Supplementary Figure 2

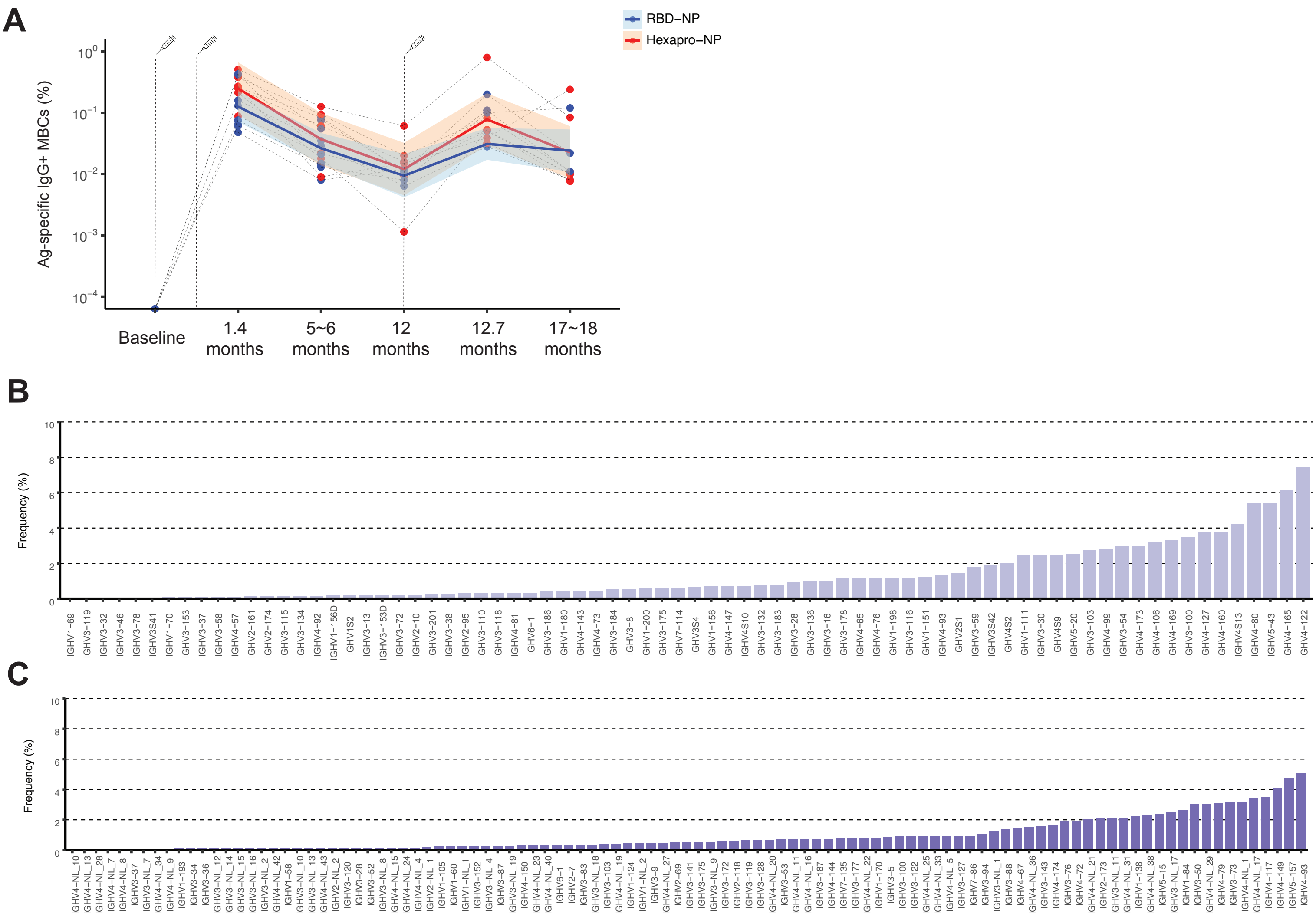


D

| Database: imgt.Macaca_mulatta.V.f.orf.p; imgt.Macaca_mulatta.D.f.orf; imgt.Macaca_mulatta.J.f.orf | | | | | | | | |
|---|---|---|---|---|---|---|---|---|
| **Query= Human_VH3-53** | | | | | **Query= Human_VH4-59** | | | |
| Length=350 | | | | | Length=350 | | | |
| Sequences producing significant alignments: | Score(Bits) | E Value | Identity | | Sequences producing significant alignments: | Score(Bits) | E Value | Identity |
| IGHV3-103*01germline gene | 377 | 1E-106 | 91.1% (267/293) | | IGHV4-122*02germline gene | 402 | 3E-114 | 93.6% (280/299) |
| IGHV3-100*02germline gene | 363 | 2E-102 | 89.9% (266/296) | | IGHV4-147*01germline gene | 397 | 8E-113 | 93.6% (277/296) |
| IGHV3S42*01germline gene | 363 | 2E-102 | 89.9% (266/296) | | IGHV4-160*01germline gene | 397 | 8E-113 | 93.6% (277/296) |

### Supplementary Figure 3

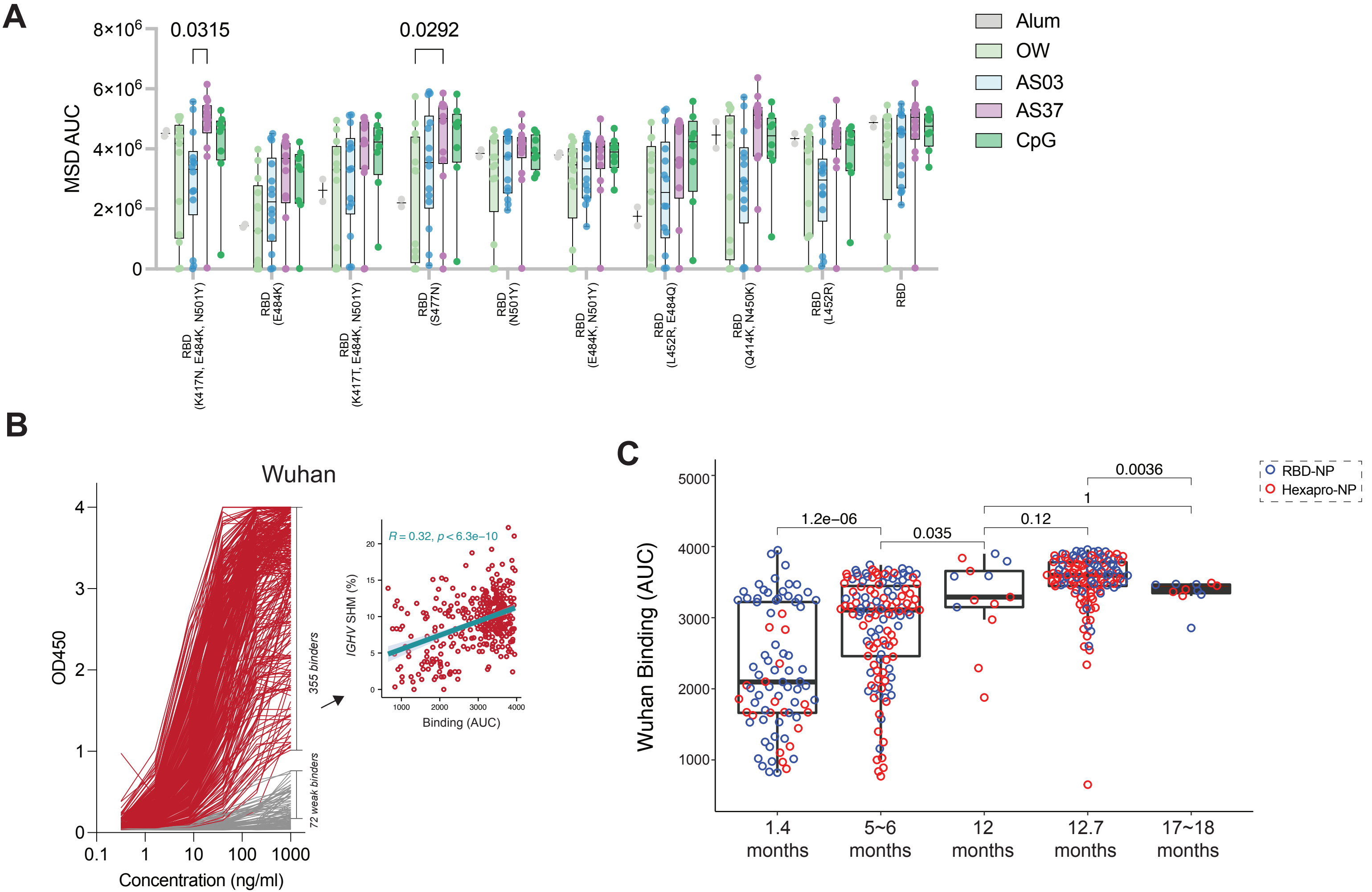

### Supplementary Figure 4

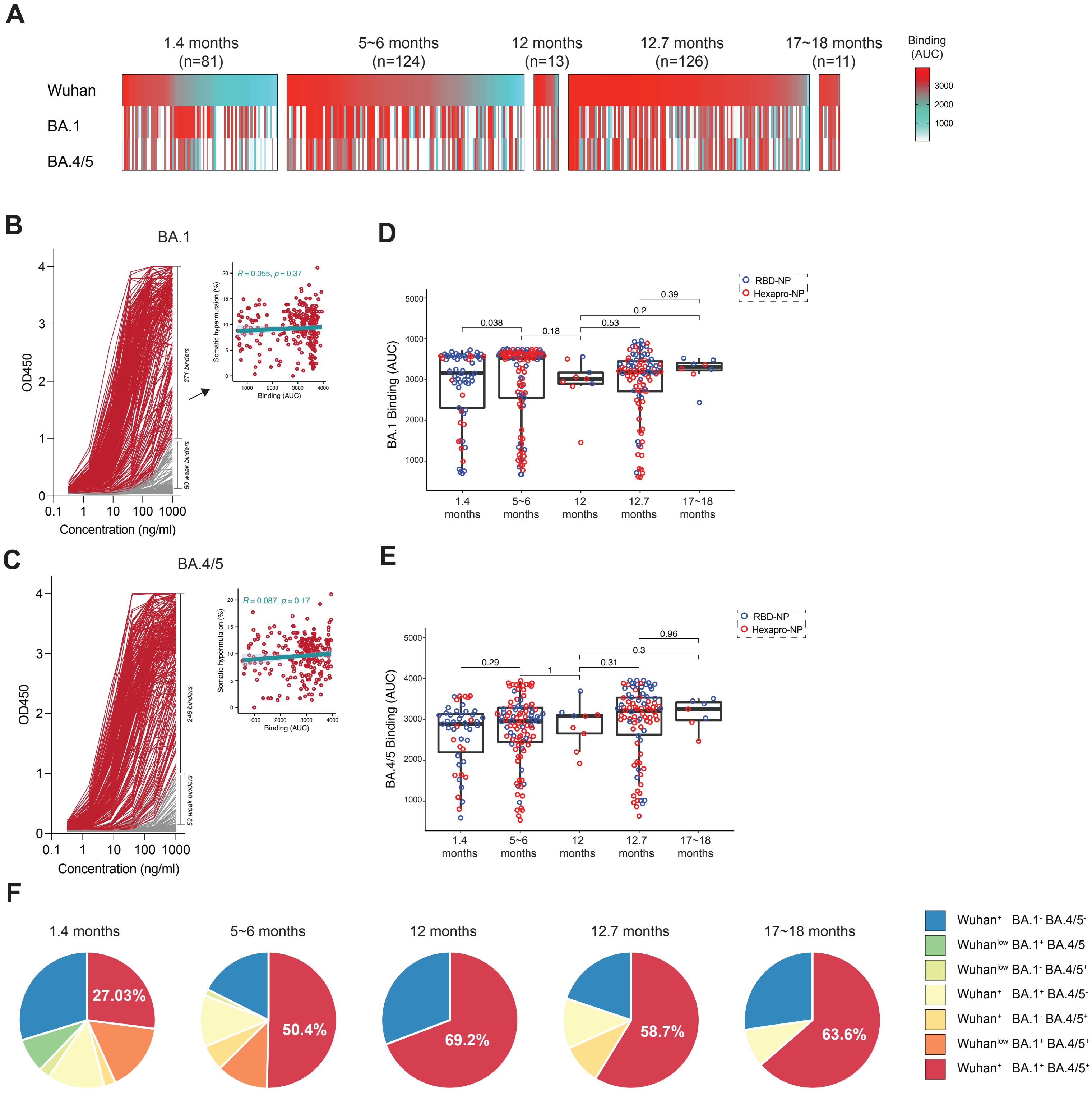

### Supplementary Figure 5

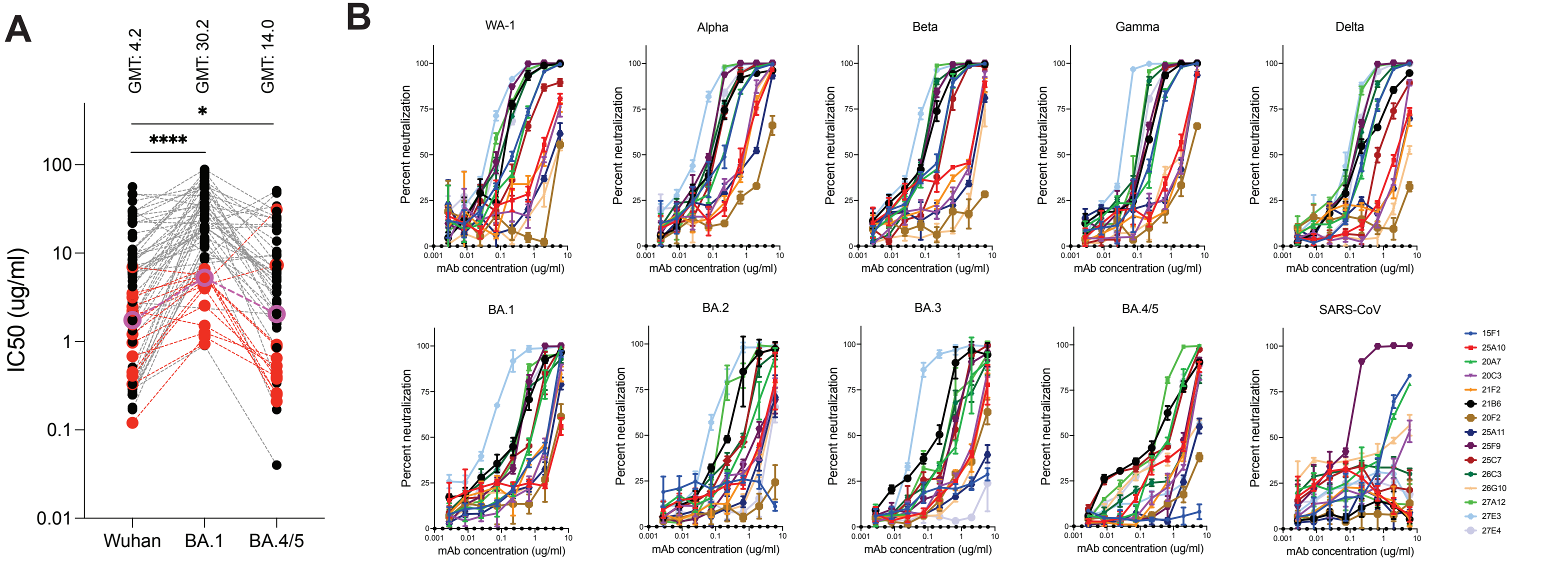


**C**

| IC50 (ng/mL) | mAb 20A7 | mAb 27A12 | mAb 27E3 | mAb 21B6 | mAb 27E4 | mAb 15F1 | mAb 25F9 |
|---|---|---|---|---|---|---|---|
| D614G | 9 | 0.89 | 0.87 | 1 | 2 | 6 | 6 |
| Omicron | 6 | 5 | 98 | 11 | 16 | 27 | 42 |
| Pangolin | 345 | 343 | | | 492 | >25000 | 6 |
| SARS-CoV | 13 | >25000 | >25000 | >25000 | 2082 | 13 | 0.85 |
| WIV1 | 2 | 695 | | | 71 | <0.32 | 3 |
| SHC014 | 18 | 9 | 12755 | >25000 | 7 | 4 | 6 |
| MERS-CoV | >25000 | >25000 | | | >25000 | >25000 | >25000 |

**D**

| | Pseudovirus neutralization (IC50 ug/ml) | | | |
|---|---|---|---|---|
| | BA.1 | BQ.1 | BQ.1.1 | XBB |
| 25F9 | 0.55056 | 6.52600 | 5.29661 | 11.92369 |
| 20A7 | 0.79407 | 5.24843 | 4.27107 | 7.65697 |
| 27A12 | 0.23749 | 0.41892 | 0.49685 | 1.26194 |
| 21B6 | 0.30181 | 11.11523 | 11.97605 | 30.00000 |

### Supplementary Figure 6

## A

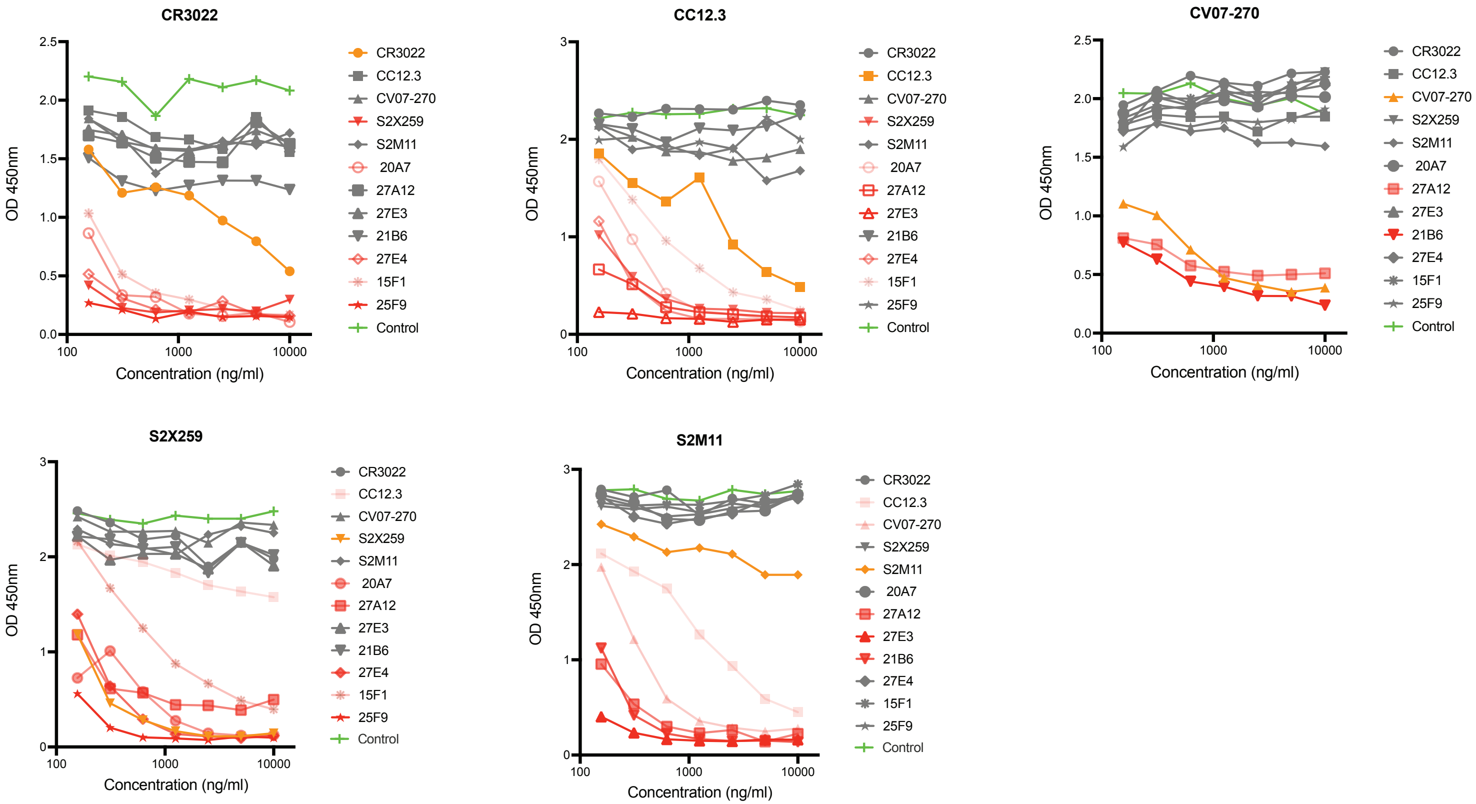


## B

| | CR3022 | CC12.3 | CV07-270 | S2X259 | S2M11 | 20A7 | 27A12 | 27E3 | 21B6 | 27E4 | 15F1 | 25F9 |
|---|---|---|---|---|---|---|---|---|---|---|---|---|
| CR3022 | 74 | 28 | 23 | 86 | 8 | 95 | 26 | 25 | 43 | 92 | 93 | 93 |
| CC12.3 | -5 | 79 | 18 | 90 | 26 | 94 | 92 | 93 | -3 | 93 | 89 | 9 |
| CV07-270 | -19 | 8 | 80 | -11 | 25 | 1 | 75 | -1 | 89 | -5 | -10 | -4 |
| S2X259 | 20 | 34 | 3 | 94 | 4 | 95 | 80 | 18 | 12 | 95 | 83 | 96 |
| S2M11 | 1 | 84 | 90 | -1 | 30 | 2 | 92 | 94 | 95 | -1 | -5 | 2 |

% Competition (color scale: 100, 50, 0)

### Supplementary Figure 7

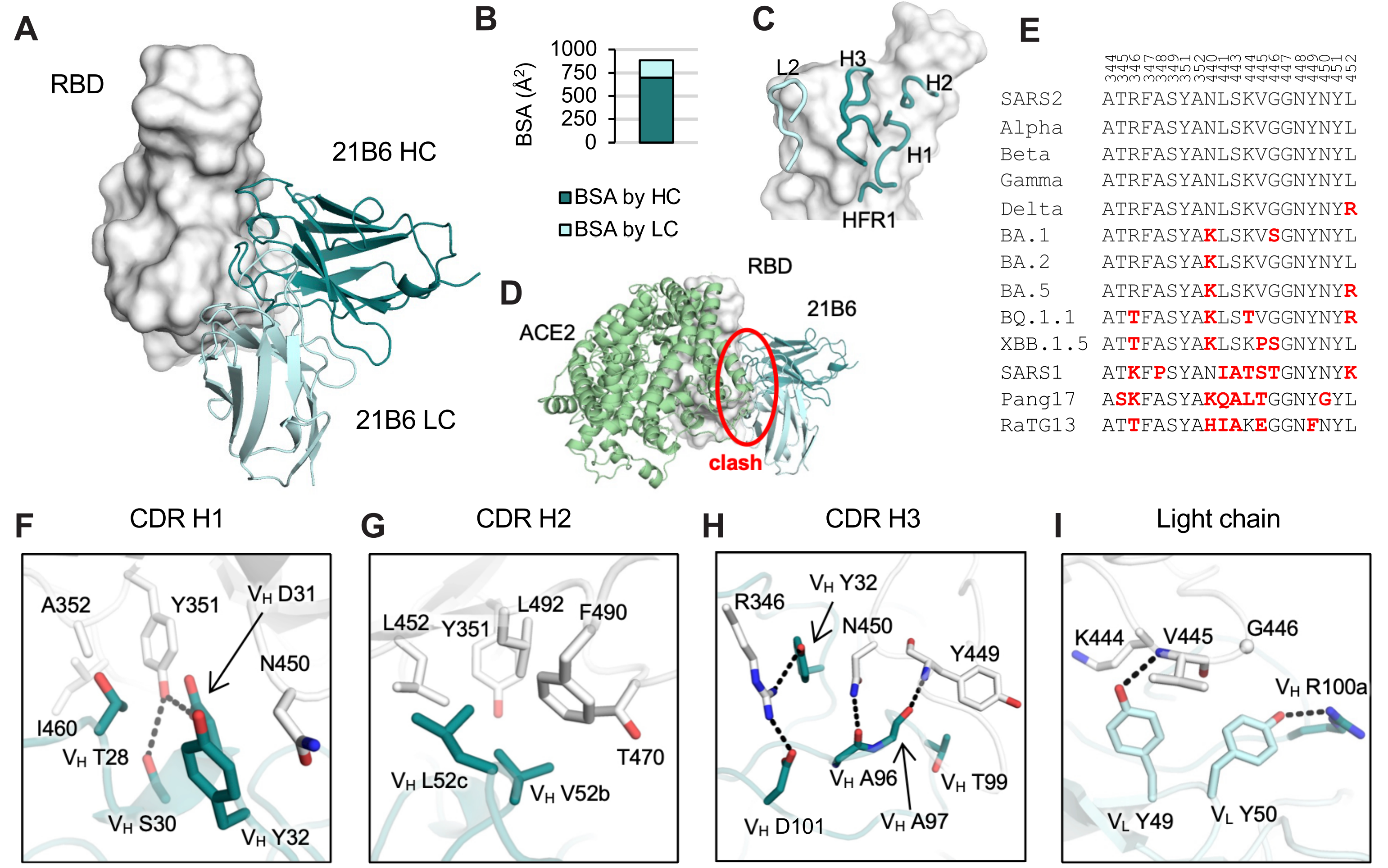

### Supplementary Figure 8

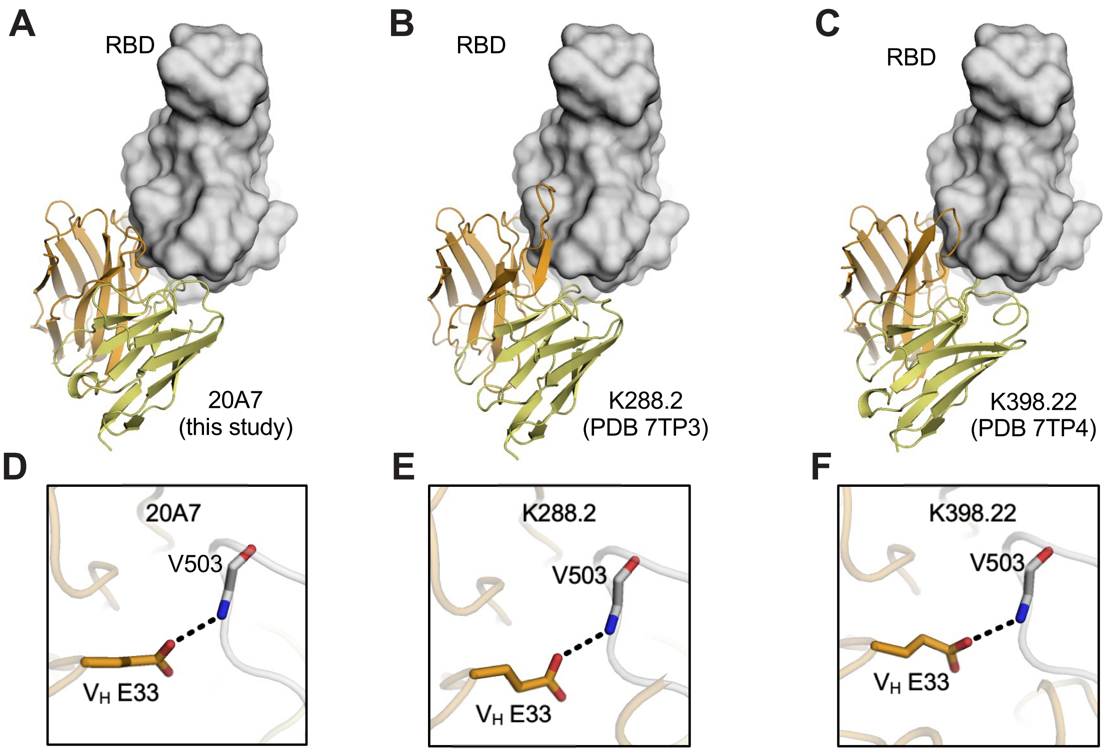


**G**

```
IGHV3-73*01_1       DVQLVESGGGLVKPGGSLRLSCVASGFTFSSYEMHWVRQAPGKGLEWVSVISESGGTTYY
IGHV3-73*01_S3636_1 DVQLVESGGGLVKPGGSLRLSCVASGFTFSSYEMHWVRQAPGKGLEWVSVISESGGTTYY
IGHV3-73*01_S6942_1 EVQLVESGGGLVKPGGSLRLSCVASGFTFSSYEMHWVRQAPGKGLEWVSVISESGGTTYY
IGHV3-73*02_1       EVQLVESGGGLVKPGGSLRLSCVASGFTFSSYVMHWVRQAPGKGLEWVSVISESGGTTYY
IGHV3-73*02_S0085_1 EVQLVESGGGLVKPGGSLRLSCVASGFTFSSYVMHWVRQAPGKGLEWVSVISESGGTTYY
IGHV3-73*02_S3937_1 DVQLVESGGGLVKPGGSLRLSCVASGFTFSSYVMHWVRQAPGKGLEWVSVISESGGTTYY
IGHV3-73*02_S4412_1 DVQLVESGGGLVKPGGSLRLSCVASGFTFSSYVMHWVRQAPGKGLEWVSVISESGGTIYY
IGHV3-73*02_S5005_1 DVQLVESGGGLVKPGGSLRLSCVASGFTFSSYVMHWVRQAPGKGLEWVSVISESGGTIYY
IGHV3-73*02_S7884_1 EVQLVESGGGLVKPGGSLRLSCVASGFTFSSYVMHWVRQAPGKGLEWVSVISESGGTIYY
IGHV3-73*02_S7955_1 EVQLVESGGGLVKPGGSLRLSCVASGFTFSSYVMHWVRQAPGKGLEWVSVISESGGTTYY
IGHV3-73*02_S8830_1 DVQLVESGGGLVKPGGSLRLSCVASGFTFSSYVMHWVRQAPGKGLEWVSVISESGGTTYY

IGHV3-73*01_1       ADSVKGRFTISRDNAKNSLFLQMNSLRAEDTAVYYCTRX
IGHV3-73*01_S3636_1 ADSVKGRFTISRDNAKNSLFLQMNSLRAEDTAVYYCTRX
IGHV3-73*01_S6942_1 ADSVKGRFTISRDNAKNSLFLQMNSLRAEDTAVYYCTRX
IGHV3-73*02_1       ADSVKGRFTISRDNAKNSLFLQMNSLRAEDTAVYYCTRX
IGHV3-73*02_S0085_1 ADSVKGRFTISRDNAKNSLFLQMNSLRAEDTAVYYCTRX
IGHV3-73*02_S3937_1 ADSVKGRFTISRDNAKNSLFLQMNSLRAEDTAVYYCTRX
IGHV3-73*02_S4412_1 ADSVKGRFTISRDNAKNSLFLQMNSLRAEDTAVYYCTRX
IGHV3-73*02_S5005_1 ADSVKGRFTISRDNAKNSLFLQMNSLRAEDTAVYYCTRX
IGHV3-73*02_S7884_1 ADSVKGRFTISRDNAKNSLFLQMNSLRAEDTAVYYCTRX
IGHV3-73*02_S7955_1 ADSVKGRFTISRDNAKNSLFLQMNSLRAEDTAVYYCTRX
IGHV3-73*02_S8830_1 ADSVKGRFTISRDNAKNSLSLQMNSLRAEDTAVYYCTRX
```

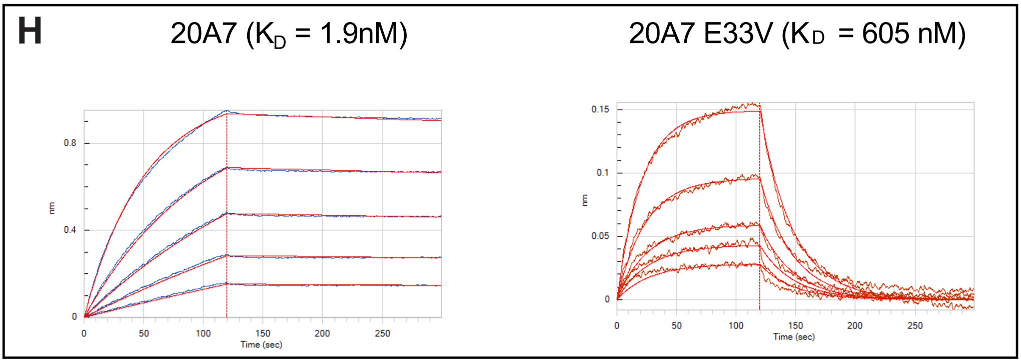

### Supplementary Figure 9

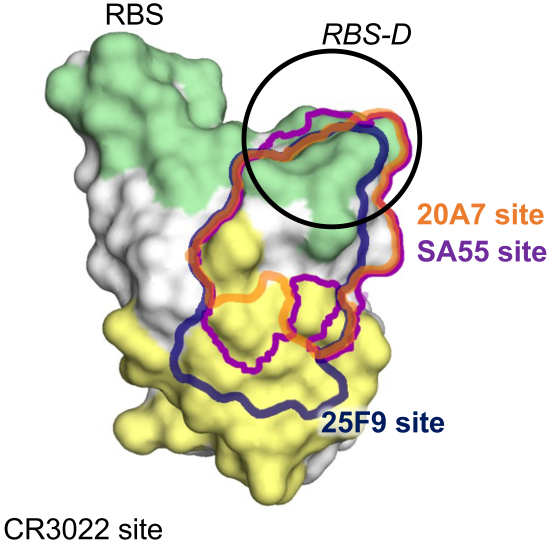
